# AN OPTONANOBODY FOR REVERSIBLE PHOTOACTIVATION OF RECOMBINANT AND NATIVE α7 NICOTINIC RECEPTORS

**DOI:** 10.64898/2026.03.10.710770

**Authors:** Maria Vangelatou, Kian Stenbolt, Sylvie Bay, Karima Medjebeur, Gabriel Ayme, Pierre Lafaye, Arnaud Blondel, Laurie Peverini, Alexandre Mourot, Pierre-Jean Corringer

**Affiliations:** Institut Pasteur, Université Paris Cité, CNRS UMR 3571, Signaling and Receptors Dynamics Unit, Paris, France; Brain Plasticity Unit, CNRS, ESPCI Paris, PSL Research University, 75005 Paris, France; Institut Pasteur, Université Paris Cité, CNRS UMR3523, Chemistry of Biomolecules, Paris 75015, France; Institut Pasteur, Université Paris Cité, CNRS UMR 3528, Antibody Engineering platform, Paris, France; Institut Pasteur, Université Paris Cité, CNRS UMR 3528, Structural Bioinformatic, Paris, France

**Author notes:** Corresponding Authors, Pierre-Jean Corringer, Alexandre Mourot.

**Keywords:** α7 nicotinic receptor, photoswitch, nanobody, photopharmacology, optonanobody, electrophysiology

## Abstract

Photopharmacology which enables the precise optical control of endogenous receptor activity, represents a powerful approach in neuroscience. However, photochromic diffusible ligands often exhibit limited subtype specificity, while alternative strategies for controlling specific receptor subtypes require genetic modification. Here, to achieve high subtype selectivity without the need of receptor engineering, we introduce a genetically independent strategy for optical control of endogenous receptors based on highly selective and chemically-defined photoswitchable nanobodies. By covalently coupling a light-sensitive azobenzene agonist to a high-affinity nanobody targeting α7 nicotinic acetylcholine receptor (nAChR), we engineered MalAzoCh-C4, an optonanobody that confers reversible, light-dependent activation of native α7 receptors. In *Xenopus* oocytes, MalAzoCh-C4 enables wavelength-controlled modulation of recombinant α7 receptors, with enhanced activity in *trans* configuration. In acute hippocampal slices, application of MalAzoCh-C4 produces robust photocontrol of endogenous α7 nAChRs in interneurons, sufficient to modulate action potential firing. This strategy combines nanobody specificity with the temporal resolution of photopharmacology, establishing optonanobodies as a platform for control of native neuronal receptors.

## INTRODUCTION

Nicotinic acetylcholine receptors (nAChRs) are members of the pentameric ligand-gated ion channels assembled from a repertoire of 17 subunits (α1-10, β1-4, γ, δ, ε) (1). Among these, α7-containing receptors have attracted much interest due to their therapeutic potential to treat neurodegenerative and psychiatric diseases, notably to develop pro-cognitive drugs (2). Homomeric α7 receptors are excitatory cationic channels with exceptionally high calcium permeability and rapid desensitization. They are expressed both at synaptic and extrasynaptic sites of neurons, as well as in non-neuronal cells such as astrocytes, microglia and immune cells. They are one of the most abundant nAChR subtypes in the central nervous system, notably in the cortex, hippocampus, hypothalamus and brain stem nuclei,where they modulate synaptic transmission, neuronal excitability, and plasticity (3).

To interrogate the contribution of nAChRs to neuronal circuit function and behavior, photochemical approaches have been developed to manipulate receptor activity with high spatiotemporal precision (4). Light offers a unique advantage in this context, as it can be delivered with millisecond temporal resolution and spatially restricted to defined cellular compartments, such as the soma or neuronal projections. Three major classes of chemical photoswitches have been developed: photochromic ligand (PCLs), photochromic tethered ligands (PTLs) and photochromic orthogonal remotely tethered ligand (PORTLs) (5-8).

PCLs are diffusible agonists or antagonists that typically incorporate an azobenzene moiety, which reversibly isomerizes between a twisted *cis* and an elongated *trans* configuration upon illumination with violet and green light, respectively (9). Early PCLs for nAChRs were based on azobenzene-grafted carbamylcholine or trimethylammonium ligands, and enabled light-dependent inhibition or activation of native muscle-type nAChRs in the electroplax of Electrophorus, with activity primarily in the *trans* configuration (10,11). These results were later confirmed on recombinant receptors (12,13). More recently, AzoCholine, a choline-based photoswitchable agonist, was reported to activate α7 nAChRs preferentially in its *trans* configuration (13,14). These studies relied on an α7/Glycine chimera, which retains the extracellular domain (ECD) and the orthosteric site of α7 but lacks desensitization (15). While PCLs have been used to control receptor activity in neuroscience (4,16), their utility is often limited by poor pharmacological specificity. This limitation is particularly relevant for simple chemical ligands such as tetramethylammonium and choline, which can act as competitive antagonists at heteromeric nAChRs (17).

PTLs and PORTLs were developed to overcome the limited pharmacological specificity of PCLs. In PTLs, photoswitchable ligands are covalently attached to a genetically-introduced cysteine residue on the target receptor, enabling cell-type and subunit-specific optical control (6,16). For nAChRs, both photochromic tethered-agonists and antagonists have been engineered, consisting of a cysteine-reactive maleimide group, an azobenzene photoswitch, and a cholinergic ligand-either acetylcholine (MAACh) or homocholine (MAHoCh). Tethering these PTLs near the orthosteric site yielded light-controlled activation (MAACh) or inhibition (MAHoCh) of α4β2 and α3β4 nAChRs (18). MAHoCh has been successfully used *in vivo* for optical manipulation in the mouse, showing that β2-containing nAChRs in the ventral tegmental area are crucial for nicotine reinforcement (19).

PORTLs were developed to circumvent the need for direct tethering near the orthosteric site and have been primarily applied to metabotropic glutamate receptors rather than nAChRs. In this approach, photoswitchable ligands are attached either to a receptor-fused SNAP-tag (20-24) or indirectly through a receptor-fused GFP that binds a nanobody-conjugated PTL (25). Because these attachment sites are distant from the orthosteric binding pocket, long and flexible polyethylene glycol linkers are required to allow the ligand to reach its target site. Although PTL and PORTL approaches provide pharmacological specificity, they require genetic manipulation of native receptors either using viral vectors (19) or through the labor-intensive engineering of a knock-in mouse line (26), which may alter receptor expression, distribution and function.

Recently, we generated a series of single chain camelid antibody fragments (nanobodies) that specifically target α7 nAChRs, but not other major brain subtypes like α4β2 and α3β4 (27). Among them, nanobody C4 was identified as a silent allosteric ligand that does not alter acetylcholine-evoked currents and binds with sub-nanomolar affinity to the top-platform of the extracellular domain (ECD) of the α7 receptor, distant from the orthosteric site and the transmembrane domain (TMD) (28). Here, we conjugated C4 to AzoCholine to generate a new class of chemically-defined photoswitchable ligands called optonanobodies. This strategy combines the temporal precision of photopharmacological tools and the molecular specificity of nanobodies, enabling photocontrol of target receptors without requiring genetic modification. We demonstrate photo-modulation of wild-type α7 receptors in *Xenopus* oocytes, as well as of native α7 nAChRs in hippocampal interneurons.

## RESULTS

### Design of a silent nanobody conjugated with a photo-controlable α7 agonist: MalAzoCh-C4

We designed a photoswitchable molecule based on AzoCholine (13) conjugated to a modified version of the α7-selective nanobody C4. AzoCholine, which consists of an azobenzene moiety fused to a choline group (13), was extended by a six-carbon chain terminating by a maleimide group for conjugation to C4, yielding MalAzoCh (Figure 1A, Figure S1). To evaluate the potential impact of this substitution on binding, we docked *cis* and *trans* MalAzoCh into the orthosteric site of the α7 cryo-EM structure 7EKP, which corresponds to an active/desensitized conformation with a capped loop C (29). Among the 40 docked poses of *cis*-MalAzoCh, 15 show a typical “agonist-like” pose with the quaternary ammonium in the aromatic box, and the maleimide chain protruding out of the pocket through the interface between the C-loop and its adjacent subunit (Figure 1B). In contrast, *trans*-MalAzoCh does not yield such poses in initial docking. We therefore generated additional structural models of *trans*-MalAzoCh bound to an α7 nAChR ECD dimer using the Boltz2 AI-based tool (30). In this context, 6 out of 16 models display an expected agonist-like pose, with the ammonium correctly positioned and loop C capped around the ligand. These docking analyses indicate that both *cis* and *trans* MalAzoCh are potential binders of active forms of the receptors, while maintaining accessibility of the maleimide group for extracellular conjugation.

**Figure 1.**
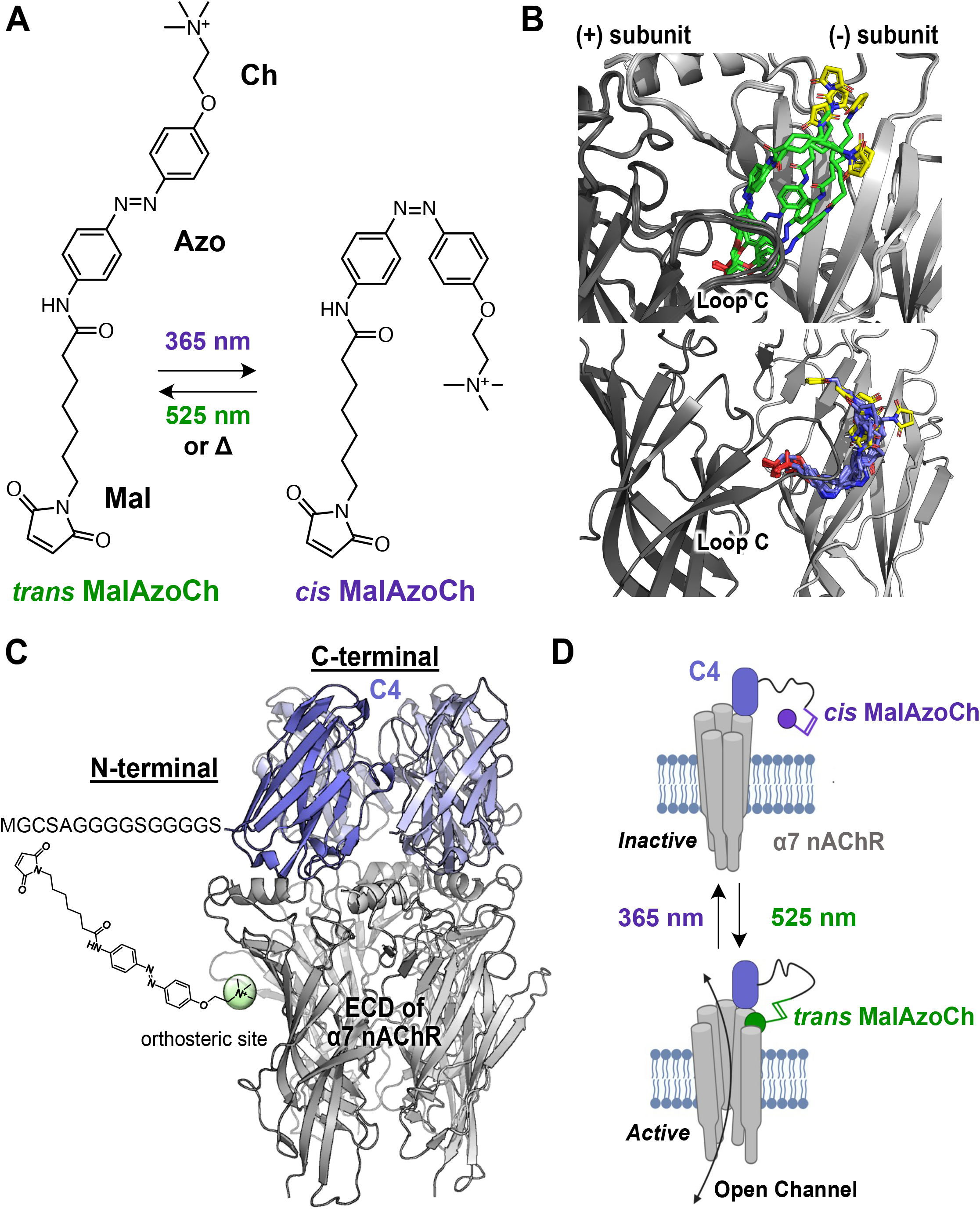
Design of MalAzoCh-C4 as a photoswitchable agonist for the optical control of the neuronal α7 nAChRs. (A) Structural formula of MalAzoCh in the *trans* and *cis* configurations upon illumination with 525 and 365 nm light, respectively. (B) Docking poses of *trans*-MalAzoCh (upper panel) and *cis*-MalAzoCh (lower panel) in the orthosteric binding site of the α7 nAChR. The maleimide moiety is shown in yellow and the quaternary ammonium of choline is shown in red. (C) Cryo-EM structure of the α7 nAChR (only the ECD is shown in grey) in complex with five C4 nanobodies, shown in violet (PDB: 8C9X). The orthosteric site is indicated with a green sphere. The Gly/Ser linker is engineered at the N-terminus of C4 where MalAzoCh is conjugated. (D) Schematic illustration of the approach of light-dependent activation of α7 nAChRs by MalAzoCh-C4.

Cryo-EM structures show that C4 binds at the top of the receptor, at the interface between two subunits and that an α7 pentamer can bind up to five C4 molecules (Figure 1C). The C4 N-terminus protrudes toward the receptor subunit interface, at an estimated straight-line distance of ∼4 nm from the orthosteric site, whereas its C-terminus is positioned further away. To allow the photoswitchable ligand to freely diffuse and reach the orthosteric site, we extended the N-terminus of C4 with a long, hydrophilic and flexible linker terminating with a distal cysteine residue (MGCSAGGGGSGGGGS) (Figure S3). The linker is designed to increase the effective local concentration of MalAzoCh, by confining it within ∼5 nm (a distance corresponding to the calculated length of the elongated linker) of the anchoring point. Under these conditions, a single MalAzoCh molecule is confined to an estimated volume of ∼500 nm^3^, corresponding to an effective local concentration in the millimolar range. This linker-based strategy, previously implemented in PORTLs, is illustrated in Figure 1D. Site-specific conjugation was performed using a maleimide/cysteine thiol-addition, yielding the chemically-defined bioconjugate MalAzoCh-C4, as demonstrated by mass spectrometry (MS) (Figure S3, Table S1).

### Non-conjugated MalAzoCh-derivative retains the photoswitchable activity of AzoCholine

To ensure that neither the linker nor the conjugation process affect the optical properties of AzoCholine, MalAzoCh was reacted with β-mercaptoethanol to generate MalAzoCh_β-mer_, (Figure 2A) bearing a succinimidyl thiolether moiety similar to that present on MalAzoCh-C4. As expected, the UV-Vis absorbance spectrum of MalAzoCh_β-mer_ recorded in the dark and under green light (525 nm) is characteristic of the *trans*-azobenzene configuration, whereas illumination with violet light (365 nm and 385nm) yields spectra of the *cis* isomer (Figure 2B) (31). High Performance Liquid chromatography (HPLC) coupled to MS analysis shows that nearly 100% of *trans* MalAzoCh_β-mer_ is found in the dark. Under green light, the *trans*/*cis* ratio is 91/9, while violet light shifts the ratio to 8/92 (365 nm) and 15/85 (385 nm). Hence, 365 nm is selected to achieve optimal *cis* photoconversion. Reversion to the *trans* state under 525 nm reaches 87% efficiency (Figure 2C). MalAzoCh_β-mer_ exhibits no sign of photo-fatigue after successive cycles of violet/green illuminations (Figure 2E). We also evaluated the stability of the *cis* isomer, which slowly relaxes back to the *trans* form in the dark with a half-life of approximately 3h (Figure 2D).

**Figure 2.**
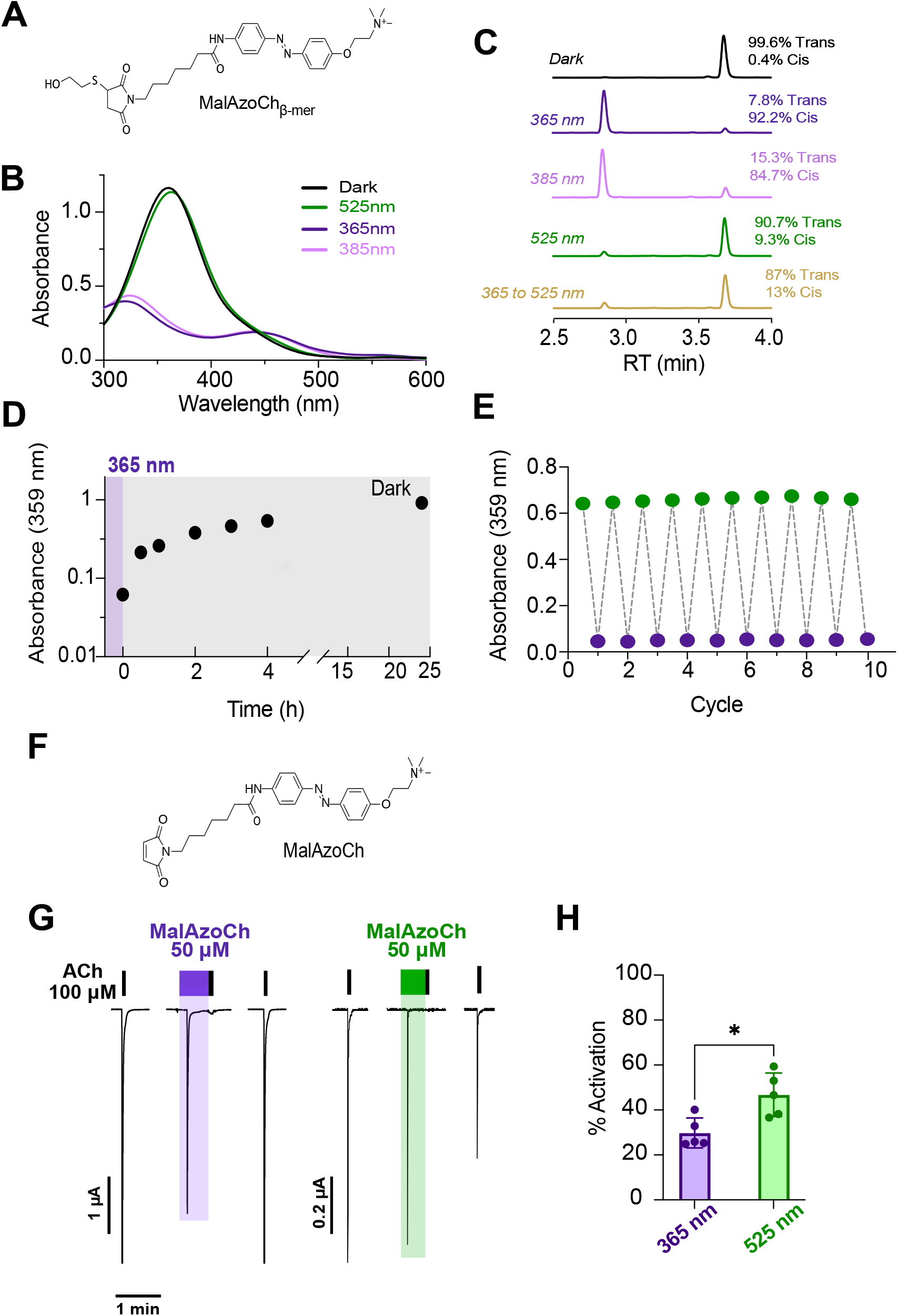
Photochemical and electrophysiological characterization of non-conjugated MalAzoCh. The photochemical characterization was conducted using MalAzoCh quenched with β-mercaptoethanol (MalAzoCh_β-mer_), while the electrophysiological characterization was conducted with free MalAzoCh. The structural formula of each compound (*trans* isomers) is presented in (A) and (F), respectively. (B) UV-Vis absorption spectra of MalAzoCh_β-mer_, recorded in the dark and after illumination at 365, 385 or 525 nm. (C) HPLC chromatograms (detection at 316 nm) of MalAzoCh_β-mer_ in the dark, after 365, 385 or 525 nm illumination, and after sequential illumination at 365 nm followed by 525 nm. *Cis* and *trans* compounds are eluted at 2.9 min and 3.7 min, respectively. (D) Relaxation kinetics of *cis*-MalAzoCh_β-mer_ in the dark, following illumination with 365 nm light (half-life t_1/2_ = 3h), as determined by absorbance at 359 nm. (E) Photoconversion profile of MalAzoCh_β-mer_ over 10 alternating cycles of illumination at 365 and 525 nm, showing no detectable photodegradation. (G) Representative current traces elicited by 100 μM ACh and 50 μM MalAzoCh, on oocytes expressing α7 nAChR WT and illuminated at 365 nm (left) or 525 nm (right). In these experiments, MalAzoCh was pre-illuminated, transferred to the perfusion system, and applied to the oocyte with simultaneous illumination (colored bar). Note the 5-10 second delay between MalAzoCh perfusion and current rise, that is potentially due to its hydrophobicity and probable partitioning into the membrane, delaying the rise of free extracellular MalAzoCh. (H) MalAzoCh -induced activation of α7 nAChRs expressed as a percentage of maximal ACh-evoked currents (n = 5 oocytes). Bars represent mean ± SD. Statistical significance was assessed using an unpaired Student’s t-test (*p < 0.05).

The activity of MalAzoCh (Figure 2F) was assessed by two-electrode voltage-clamp electrophysiology (TEVC) in *Xenopus* oocytes, under green or violet illumination. For each oocyte, an ACh pre-perfusion at 100 μM (near the EC_50_, see Figure S5C and Table S2) was applied to evaluate the oocyte response, followed by a 3-minute rinsing period to allow the receptor to recover from desensitization. MalAzoCh was then perfused for 30 seconds, immediately followed by an ACh post-perfusion application to quantify the extent of desensitization. ACh pre-perfusion elicits robust currents with rapid onset and offset that are characteristic of α7 fast activation and desensitization. Likewise, application of 50 μM MalAzoCh elicits activation followed by desensitization, with time-courses similar to those evoked by ACh. Peak current amplitudes reach 30% ± 6.6 and 47% ± 9.7 of ACh maximal currents under violet and green illumination, respectively. Subsequent ACh perfusions fail to evoke significant current, suggesting complete desensitization (Figure 2G, H). These results show that MalAzoCh is an agonist of α7 that is more active in the *trans* than in the *cis* configuration.

### MalAzoCh-C4 activates and desensitizes α7 nAChRs, with higher activity in *trans*

The mutated C4 was produced and purified as described previously by (27) and exhibits no intrinsic modulatory activity in TEVC (Figure S5A, B). Following conjugation with MalAzoCh (Figure S3 and S4), MalAzoCh-C4 acts as an agonist since it elicits a small but significant current under green light at high concentration (5 μM; 7% of maximal ACh-evoked currents; Figure 3B, D). These currents display slower onset and offset kinetics compared to ACh-evoked responses. Subsequent ACh post-perfusion elicits no significant current, suggesting complete desensitization. Concentration-response analysis reveals detectable currents starting at 600 nM MalAzoCh-C4, while ACh post-perfusion currents start to decrease at 300 nM and follow a sigmoidal curve with IC_50_= 574 ± 92.2 nM and nH= 2.13 ± 0.52. Under violet light, MalAzoCh-C4 currents are of smaller amplitudes and are detectable only at 3 μM, while the ACh post-perfusion curve is shifted approximately two-fold to the right (IC_50_= 1197 ± 222.7 nM and nH= 11.47 ± 10.12) (Figure 3A, C). Kinetic analysis also indicated a faster rise-time for the currents in green compared to violet light (Figure S7A, B, Table S3). Of note, MalAzoCh-C4 produces no detectable response at α4β2 receptors (Figure S6), consistent with high specificity of C4 for α7 nAChRs (27, 28). However, it also activated heteromeric α7β2 receptors (Figure S8), suggesting that its activity is not restricted to homomeric α7 assemblies.

**Figure 3.**
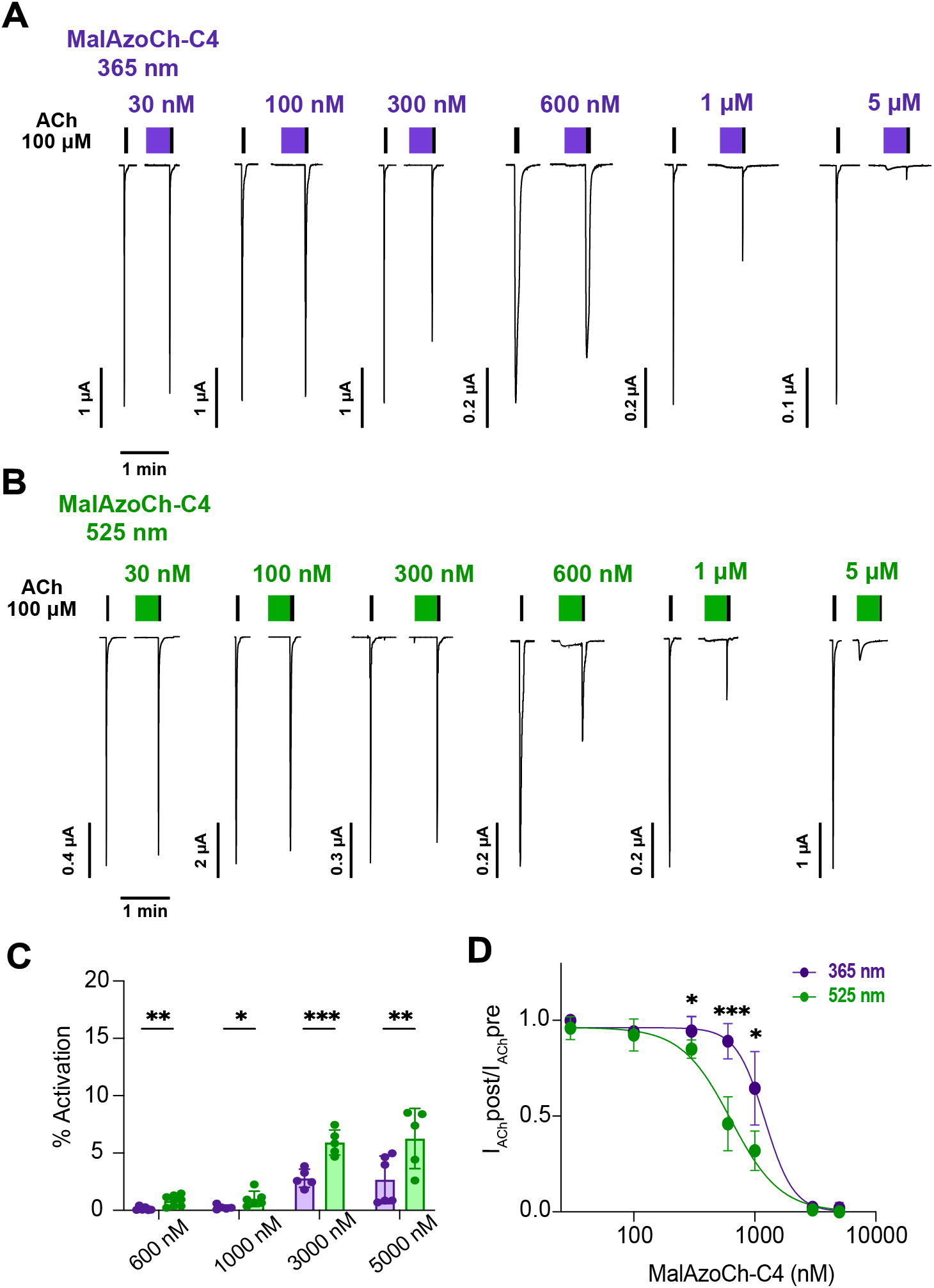
Electrophysiological analysis of MalAzoCh-C4 on α7 WT. (A) Representative current traces elicited by successive applications of 100 μM ACh, a 3-min wash, a 30 s application of MalAzoCh-C4 under illumination at 365 nm, and a 100 μM post-perfusion ACh. Each trace corresponds to a different oocyte. (B) Same as in A., for illumination at 525 nm. (C) Concentration-response bar graph for MalAzoCh-C4-elicited activation (n = 5-8). (D) Concentration-response curves of MalAzoCh-C4-elicited desensitization, quantified as I_AChpost_/I_AChpre_, under 525 nm (green curve, IC_50_= 574 ± 92.2 nM, nH= 2.13 ± 0.52, n = 5-8) and 365 nm illumination (violet curve, IC_50_= 1197 ± 222.7 nM, nH= 11.47 ± 10.12; n = 5-6). Statistical differences between 525 nm and 365 nm conditions for desensitization and activation were assessed using an unpaired Student’s t-test. *, p < 0.05; **, p < 0.01; ***, p < 0.001.

Together, these results demonstrate that MalAzoCh-C4 elicits weak but significant maximal currents under both illumination conditions, while prolonged application strongly inhibits subsequent ACh-evoked currents, most likely due to desensitization. It is noteworthy that, like most nanobodies, C4 binds slowly to the receptor, as previously shown by surface plasmon resonance on purified α7 nAChRs (28). In line with this, the rise time of MalA-zoCh-C4-evoked currents range from 2 to 20 s, which is substantially slower than the intrinsic desensitization kinetics of α7 nAChRs, occurring on the order of 700 ms (Table S3). Assuming that MalAzoCh-C4 binding elicits fast receptor activation and desensitization, we suggest that by the time peak currents are reached, a substantial fraction of receptors has already undergone the activation and desensitized transition, leading to an underestimation of the actual fraction of receptors transiently activated.

### MalAzoCh-C4 robustly activates and desensitizes α7 L247T, with higher activity in *trans*

To further assess the impact of desensitization on peak current amplitudes, we used the α7 L247T mutant, which displays markedly slowed desensitization kinetics. This mutation disrupts a central hydrophobic ring in the channel and confers a gain-of-function phenotype, characterized by a 20-fold decrease in the ACh EC_50_ (Figure S5C and Table S2). Under green light, application of 1μM MalA-zoCh-C4 elicits robust currents, reaching 80% of the response elicited by a near maximal ACh concentration (30 μM). Subsequent ACh post-perfusion fails to produce additional activation (Figure 4A, B). Both activation and desensitization are detected at MalAzoCh-C4 concentrations as low as 1 nM and increase in a concentration-dependent manner, with desensitization following a sigmoid curve with an IC_50_ of 26.1 ± 21.1 nM, nH= 0.83 ± 0.36. Under violet light, we observe a marked rightward shift of the activation and desensitization curves, which is 20-fold for desensitization (IC_50_= 606.5 ± 320.7 nM, nH= 5.45 ± 6.01) (Figure 4C, D). Finally, MalAzoCh-C4-elicited currents are inhibited by two α7 antagonists, the selective competitive antagonist methyllycaconitine (MLA) and the channel blocker memantine, confirming that the responses are mediated by α7 nAChRs (Figure S9). Overall, these results further suggest that the weak peak current amplitudes observed in WT α7 are, at least in part, attributable to fast desensitization. MalAzoCh-C4 is a more active agonist under green than violet light, a feature that is particularly pronounced in the slow-desensitizing α7 L247T mutant.

**Figure 4.**
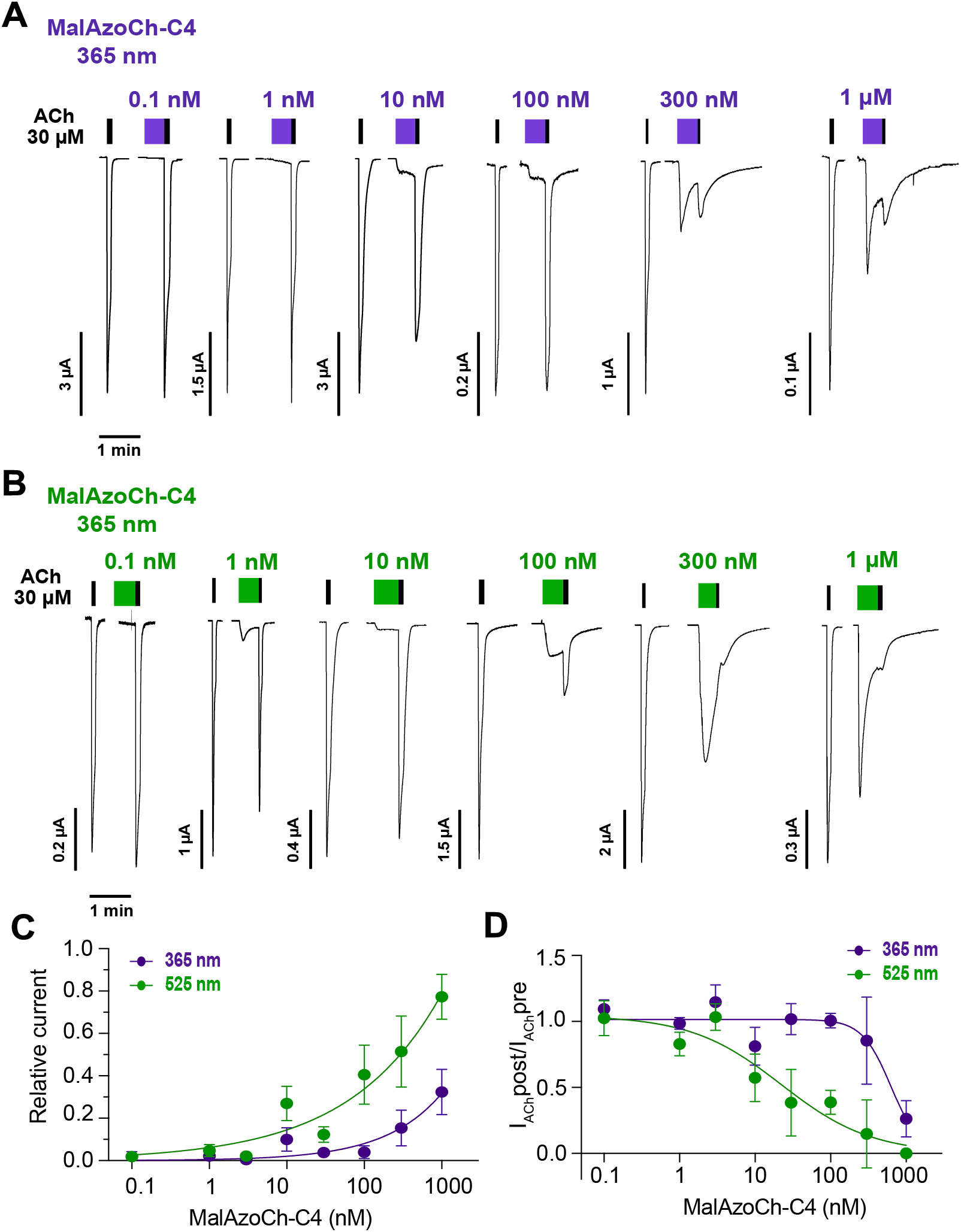
Electrophysiological analysis of MalAzoCh-C4 on α7 L247T. (A) Representative current traces elicited by successive applications of 30 μM ACh, a 3-min wash, a 30 s application of MalAzoCh-C4 under illumination at 365 nm, and a 30 μM post-perfusion ACh. Each trace corresponds to a different oocyte. (B) Same as in A., for illumination at 525 nm. (C) Concentration-response curves of MalAzoCh-C4-elicited activation, quantified as I_MalAzoCh-C4_/I_AChpre_, under 525 nm (green curve) and 365 nm (violet curve) illumination (n = 5-6). (D) Concentration-response curves of MalAzoCh-C4-elicited desensitization, evaluated as I_AChpost_/I_AChpre_, under 525 nm (green curve, IC_50_= 26.1 ± 21.1 nM, nH= 0.83 ± 0.36, n = 5-6) and 365 nm (violet curve, IC_50_= 606.5 ± 320.7 nM, nH= 5.45 ± 6.01, n = 5-6) illumination.

Residual activity in violet light likely reflects the remaining fraction of *trans* MalAzoCh-C4 (around 10% at photo-stationary state), and possibly a weak intrinsic activity of the *cis* isomer, although the latter cannot be directly resolved from our experiments.

Of note, we also conjugated AzoCholine to the C-terminus of C4 using a longer 16 amino-acid linker to compensate for the greater distance to the orthosteric site (Figure S3, S4). This C4-MalAzoCh construct likewise functions as an α7 agonist (Figure S10), showing that the strategy can be applied to both extremities of the nanobody scaffold.

### Light-controlled activation by MalAzoCh-C4 in*Xenopus* oocytes

The different efficacies of MalAzoCh-C4 in green and violet light suggest that this nanobody could be used to photocontrol α7 nAChRs, i.e. turn receptors on and off reversibly. However, reversible switching is challenged by the rapid desensitization of α7 receptors, particularly at high nanobody concentrations.

We first tested reversible photomodulation in the slow-desensitizing α7 L247T mutant. We selected 10 nM MalA-zoCh-C4, a concentration eliciting sizable currents while limiting desensitization to around 50% under green light. During prolonged perfusion under violet light, currents reach a steady state level that increases ∼3-fold upon switching to green light, consistent with the dose-response relationship (Figure 5A). A second illumination cycle produces a return to the baseline under violet light and a ∼2-fold increase under green light, and this bidirectional photomodulation can be reliably reproduced over multiple cycles. Control experiments show that no significant photomodulation occurs in oocytes expressing α7 L247T during ACh perfusion alone (Figure S11).

**Figure 5.**
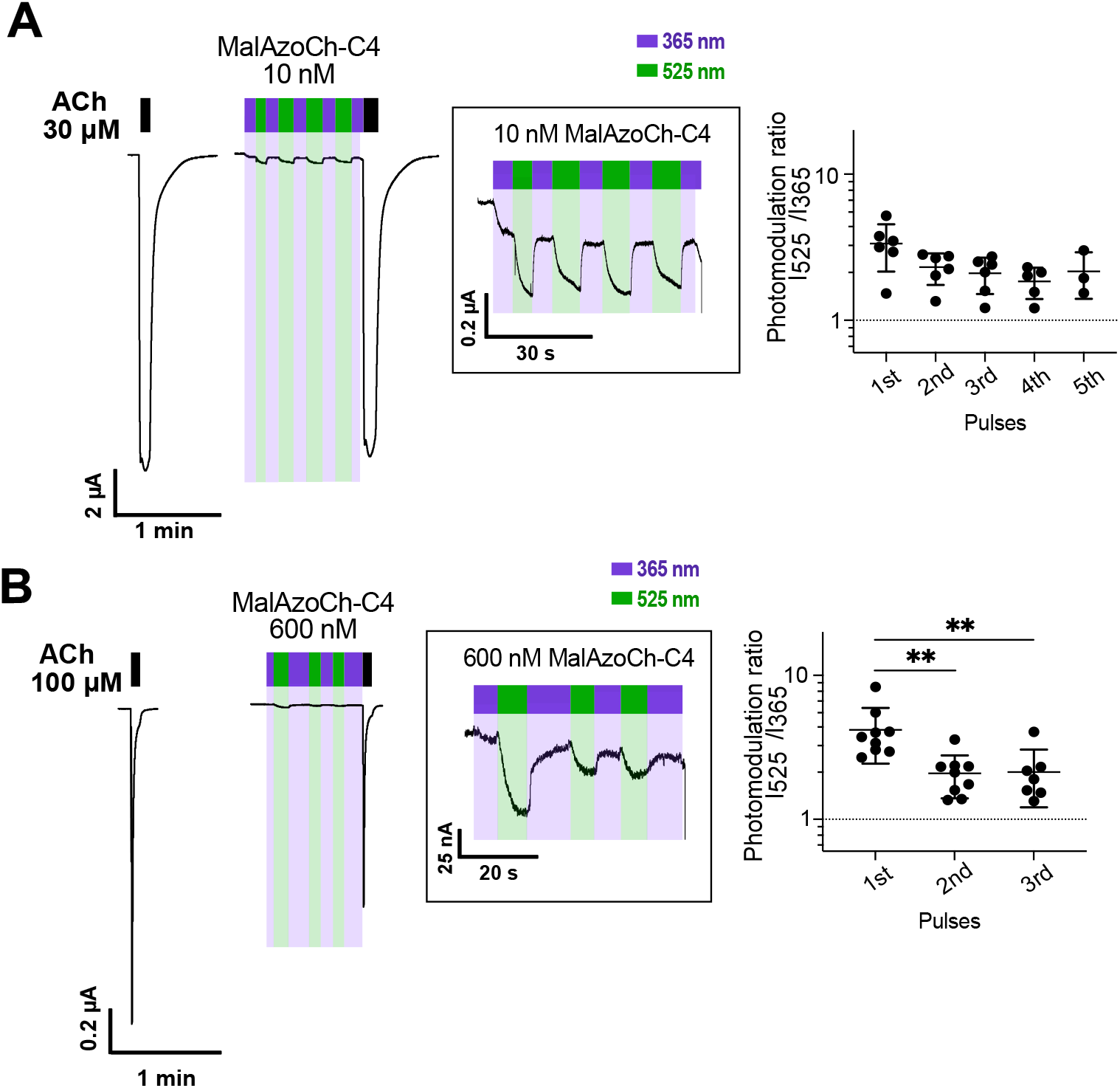
Reversible photomodulation of α7 in oocytes. Representative current traces from oocytes expressing (A) α7 L247T during prolonged application of 10 nM MalAzoCh-C4 and (B) α7 WT during a prolonged application of 600 nM MalAzoCh-C4, under alternating illumination at 365 nm (violet bars) and 525 nm (green bars). A magnified view of the photomodulation cycles is provided on the right and the photomodulation ratio (I_525_/I_365_) calculated for each 525 nm illumination pulse. For each oocyte (n = 6), at least three green light pulses were applied. Ordinary one-way ANOVA revealed no significant difference in photomodulation values between pulses (ns, p > 0.05).

For WT α7 receptors, we used 600 nM of MalAzoCh-C4, a concentration that produces significant activation while exerting minimal desensitization under violet light (Figure 3C). Alternating violet/green illumination cycles results in a reproducible ∼2-fold modulation of current amplitude, although responses are smaller than those observed for the L247T mutant (Figure 5B).

Together, these results demonstrate that MalAzoCh-C4 enables reversible photomodulation of α7 nAChRs under conditions that minimize desensitization. Although the currents amplitudes are small in comparison with maximal ACh-evoked responses, particularly for WT α7, the photomodulation is reproducible and in principle suitable for probing native receptor functions.

### Light-controlled activation of native α7 receptors in hippocampal interneurons

To investigate whether MalAzoCh-C4 enables photocontrol of native receptors, we performed whole-cell patchclamp recordings in acute hippocampal slices, focusing on GABAergic interneurons located in the stratum radiatum near the stratum molecular, a region enriched in α7-containing nAChRs (32-34). A brief (200 ms) local puff application of 3 μM MalAzoCh-C4 was delivered directly onto the soma of recorded neurons, to minimize receptor desensitization and increase peak currents as compared to the relatively slow perfusion in TEVC (Figure 6A). Under these conditions, MalAzoCh-C4 reliably elicits inward currents that are blocked by MLA, confirming they are mediated by α7-containing nAChRs (Figure 6B). The amplitude of the evoked currents is 13% of those evoked by 3 mM ACh (16.72+/-2.57 pA for MalAzoCh-C4 compared to 126.1 +/-16.56 pA for ACh, Figure 6C), which contrasts with the results obtained in oocytes, where green light-induced currents represented only ∼1% of maximal ACh responses. Importantly, non-conjugated MalAzoCh (3 μM) fails to elicit significant current, indicating conjugation to C4 is required to increase binding affinity and achieve activation of α7 receptors (Figure S12).

**Figure 6.**
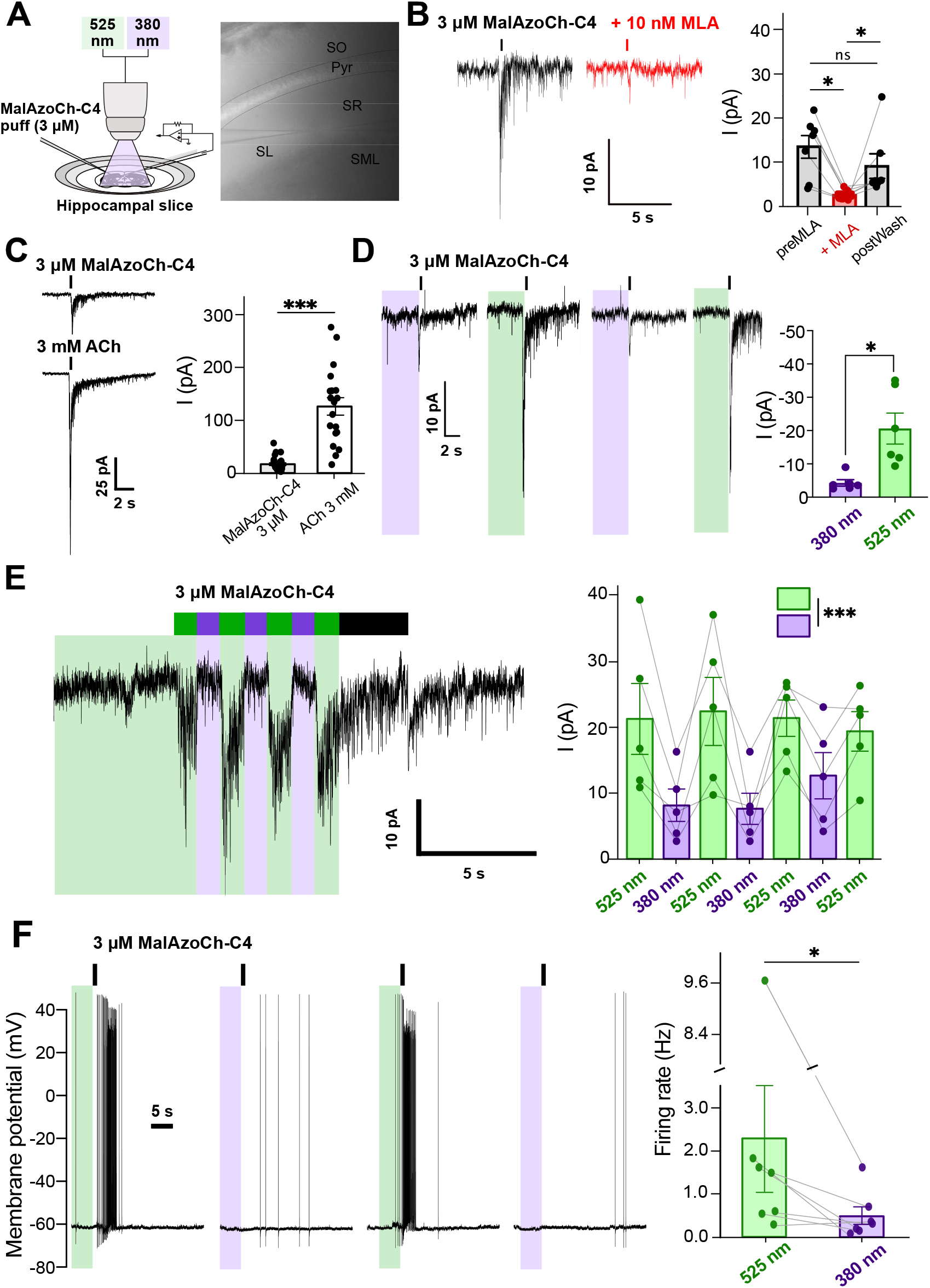
Light-dependent modulation of native α7 nAChRs in hippocampal interneurons. (A) Left, schematic representation of the patch-clamp configuration. A puff pipette delivers 3 µM MalAzoCh-C4 and illumination is provided by LED lamps emitting either 380 nm (violet) or 525 nm (green) light. Right, low-magnification (4×) image of a hippocampal CA1 slice showing the placement of the patch pipette and puff pipette in the stratum radiatum (SR), near the border with the stratum molecular (SLM), a region enriched in interneurons expressing α7-containing nAChRs. Anatomical layers are indicated: SO, stratum oriens; Pyr, stratum pyramidale; SLM, stratum molecular (right panel). (B) Left, representative voltage-clamp traces showing inward currents evoked by local application of 3 µM MalAzoCh-C4 (puff 200 ms, black bar) under control conditions (top trace, black) and in the presence of 10 nM methyllycaconitine (MLA), a selective α7 nAChR antagonist (bottom trace, red). Right, quantification of MalAzoCh-C4-evoked current before, during and after MLA application (n = 7 neurons from 5 mice, Friedman test with Wilcoxon signed-rank post hoc test). (C) Representative current traces (left) and average current (right) evoked by puff applications (200 ms) of MalAzoCh-C4 (3 µM) or acetylcholine (3 mM). Currents evoked by MalAzoCh-C4 correspond to 13.26% of the current evoked by saturating ACh (n = 26 neurons from 12 mice for MalAzoCh-C4, 19 neurons from 5 mice for ACh, p = 7.51e-10, Wilcoxon rank-sum test) (D) Left, representative current traces showing responses to 3 µM MalAzoCh-C4 under 380 nm (violet, left) and 525 nm (green, right) illumination. Light was applied for 5 seconds on the pipette containing MalAzoch-C4 prior to the puff application (indicated by color bars). Right, application of 3 µM MalAzoCh-C4 produced larger responses under green light (525 nm: 20.60 ± 11.41 pA) compared to violet light (380 nm: 4.28 ± 2.39 pA; n = 9 cells from 5 mice, Wilcoxon signed rank exact test). (E) Left, representative trace showing the response to a prolonged (10 s) puff application of 3 µM MalAzoCh-C4 while alternating illumination (initial 5 s green light, followed by three cycles of 380 and 525 nm light). Right, current amplitude increased with green light and decreased upon switching to violet light (n= 5 cells from 2 mice; t-test with holm correction). (F) Left, representative current-clamp traces showing membrane potential responses to puff applications of 3 µM MalAzoCh-C4 (1 s) under 380 nm (violet) or 525 nm (green) illumination. Light was applied 5 s prior to the puff. Right, neurons exhibited a marked increase in action potential firing under green light, whereas firing was greatly reduced or absent under violet light, (n = 7 cells from 6 mice, Paired Wilcoxon signed-rank test). For all panels, Bars represent mean± SEM. * p < 0.05; ** p < 0.01.

To assess light dependence, slices were pre-illuminated for 5 seconds before puff application. Robust inward currents are observed under green light, whereas responses are minimal under violet light (Figure 6D). These results are consistent with TEVC recordings but exhibit a more pronounced photoswitching amplitude. We next examined photomodulation during sustained perfusion of 3 μM MalAzoCh-C4 while alternating illumination conditions. Application under green light induces a robust current, which is suppressed more than 4-fold upon switching to violet illumination. This photomodulation pattern is reproducible across multiple cycles (Figure 6E). The larger current amplitudes and enhanced photocontrol observed in native neurons compared to oocytes may reflect faster ligand delivery, higher effective light intensity at the focal plane, and/or intrinsic differences between native and recombinant receptors (see Discussion).

Finally, we explored the effect of MalAzoCh-C4 on neuronal excitability using current-clamp recordings. Application of 3 μM MalAzoCh-C4 under green light markedly increases action potential firing rate, while the effect is strongly reduced under violet light (Figure 6F). This photoreversible modulation of excitability highlights the ability of MalAzoCh-C4 to exert light-dependent control over neuronal activity via native α7-containing nAChRs. Collectively, these results demonstrate that MalAzoCh-C4 enables reversible, optical control of native α7-containing nicotinic receptors in brain tissue and provide an efficient tool for studying their functional roles in neuronal circuits.

## DISCUSSION

Here we developed a new class of photochromic ligands, optonanobodies. By conjugating a photoswitchable ligand to a high-affinity subtype-specific nanobody, we combine molecular targeting with photopharmacological precision, enabling wavelength-dependent modulation of native receptor function without genetic modification. Using α7 nicotinic acetylcholine receptors (nAChRs) as a model system, we demonstrate that this approach supports robust and reversible photocontrol in intact brain tissue and modulation of neuronal excitability.

We previously generated a series of α7-specific nanobodies that bind a common epitope on the receptor top platform. Among these, C4 acts as a silent allosteric ligand (SAL), E3 acts as a positive allosteric modulator (PAM) (27), while F1 and E6 act as negative allosteric modulators (NAM) (35). By conjugating the photoswitchable agonist AzoCholine to C4, we transformed a silent binder into an optically addressable agonist, thereby expanding this nanobody toolbox toward direct and reversible receptor activation. Given the therapeutic relevance of α7 nAChRs, numerous small synthetic agonists and PAMs have been developed (36). However, clinical trials for Alzheimer’s, Parkinson’s diseases and schizophrenia have shown equivocal results and/or adverse effects, often attributed to insufficient subtype selectivity or off-target activity (37). To date, none has received regulatory approval for clinical use. By contrast, nanobody-based ligands offer high subunit specificity through defined epitope recognition, and recent work further highlights their capacity to modulate receptor function *in vivo* (38). Our approach builds on this principle by converting a targeting nanobody into a photoswitchable agonist capable of controlling native receptor function with high spatio-temporal and pharmacological precision.

Our strategy to design MalAzoCh-C4 was to conjugate AzoCholine to the N-terminus of C4 via a 13 amino acid flexible linker. This configuration allows AzoCholine to explore the vicinity of the agonist binding site while remaining spatially confined near the receptor, thereby increasing its effective local concentration into the millimolar range. Conceptually, this strategy differs fundamentally from the classical PTL approach, in which the photoswitchable ligand is directly tethered to the receptor and acts as a rigid, articulated arm that positions the ligand within or outside of the orthosteric site in a light-dependent manner (16). It also differs from nanobodies fused to LOV domains, that allow light-controllable binding of nanobodies to their target (39). Instead, our system more closely resembles the PORTL approach, yet with the fundamental advantage that it does not require genetic modification of the receptor (e.g., SNAP-tag or GFP fusion). Optical control is achieved through nanobody targeting rather than receptor engineering. This modular and versatile strategy also opens the door to rational re-engineering: the choline headgroup could be substituted by virtually any orthosteric or allosteric ligand, linker length and flexibility can be tuned to optimize effective concentrations and efficacy, and multivalent nanobody formats could increase avidity and prolong receptor binding (27). Although demonstrated here with a cholinergic agonist and α7 nAChRs, this strategy is conceptually transferable to other ion channels, G protein–coupled receptors, and membrane proteins for which high-affinity binders are available.

A central aim of the study was to achieve reversible photocontrol of WT α7 receptor activity. In oocytes, photocontrol is constrained by rapid desensitization. Optimal conditions would require a concentration of MalAzoCh-C4 that elicits substantial currents in green light, while inducing minimal desensitization in violet light, thereby allowing receptor recovery. In practice, this concentration window proved to be very narrow: at 600 nM MalAzoCh-C4, green light elicited currents that corresponded to only∼1% of maximal ACh responses. In contrast, in hippocampal interneurons, a brief puff of 3 μM MalAzoCh-C4 produced robust and repeatable photomodulation, with green-light-evoked currents corresponding to 13% of maximal ACh responses. These currents are sufficient to drive action potential firing, demonstrating physiologically relevant optical control of neuronal excitability. The enhanced photocontrol observed in native tissue likely reflects multiple factors. Rapid local ligand delivery in patchclamp recordings reduces the overlap between binding and desensitization. Optical stimulation in slice recordings is characterized by higher photon density and is confined to the neuronal soma, resulting in more efficient photo-conversion. Additionally, intrinsic differences between native and recombinant receptors may contribute: CA1 interneurons express both homomeric α7 and heteromeric α7β2 nAChRs (40,41), and α7β2 nAChRs desensitize more slowly than homomeric α7 in both oocytes and neurons (42,43). Because MalAzoCh-C4 also activates α7β2 nAChRs (Figure S8), the presence of these heteromeric receptors in interneurons may contribute to the enhanced photocontrol observed in native tissue.

MalAzoCh-C4 enables reversible optical control of native α7 receptor activity and neuronal excitability. This optonanobody thus constitutes a powerful tool to dissect α7 nAChR function in hippocampal circuits with high spatio-temporal resolution, to control important processes such as long-term potentiation and depression of synaptic activity, or oscillatory activity such as theta rhythms (44), in relation to high order cognitive function and pathologies associated with α7 nAChRs. In this context, several limitations of MalAzoCh-C4 can be foreseen, although they can be overcome by further molecular engineering. In brain slice, MalAzoCh-C4 elicits only partial receptor activation and dissociates relatively rapidly, requiring continuous perfusion to maintain photocontrol. These limitations could potentially be overcome in the future through the engineering of C4 conjugates bearing multiple MalAzoCh moieties, and/or bivalent C4/C4 constructs that should display much slower dissociation kinetics. *In vivo*, MalA-zoCh-C4 will likely need to be administered directly into the brain region of interest, as previously demonstrated for caged ligands with optofluidic devices (45), or possibly via intraventricular injection, since nanobodies are known to diffuse within the brain (46). Intravenous injection could also be considered since nanobodies can cross the blood brain barrier (BBB) (38). In addition, because MalA-zoCh-C4 exhibits higher activity in green/darkness than under violet illumination, this property may limit its use following intraventricular or systemic administration, where controlling the basal receptor activation prior to illumination could be more challenging. However, photoswitchable ligands can be designed to display higher activity in the *cis* configuration (47) or to possess thermodynamically stable *cis* isomers (48), thereby minimizing dark activity and improving applicability.

Beyond α7 receptors, optonanobodies could enable selective optical control of membrane proteins for which subtype-selective small-molecule photoligands are unavailable. More broadly, this work illustrates that spatial confinement by molecular recognition can be harnessed to transform weak photoswitchable ligands into effective and reversible modulators of native receptors. By separating optical control from genetic manipulation, optonanobodies expand the scope of photopharmacology to endogenous targets in intact tissue.

## MATERIALS AND METHODS

### Models of MalAzoCh in complex with α7-nAChR

Two complementary approaches were employed: coconstruction using a deep-learning model approach, and molecular docking onto selected receptor structure from the PDB.

For the co-construction of α7-nAChR and MalAzoCh, Boltz2 was used, an open-source AI model developed under MIT license to democratize alphafold3 level accuracy of structural 3D complex predictions (30). Sixteen copies of a model composed of two extracellular α7 nAChR chains and a copy of MalAzoCh were built using the MSA server (49).

For the second approach, MalAzoCh was docked onto α7-nAChR structure 7EKP (at the interface between chains A and B) with Autodock Vina 1.1.2 (50). Residues 54, 79, 141, 183, 186 of chain B were selected as flexible, as they could prevent the tail of the molecule from reaching the zone where the nanobody binds. The docking box was adjusted to comprise the orthosteric site and the flexible residues (center: 146.679, 135.239, 85.467; size: 18.673, 20.705, 32.695). The ligand was prepared as in (51), yielding one *trans* form and 2 *cis*, “P” and “M” slightly helical forms avoiding phenyl rings steric hindrance. The receptor was protonated with ChimeraX 1.9. Up to twenty poses were recorded after docking with an exhaustiveness of 96. Of the 20 *trans* + 40 *cis* poses obtained, those presenting their choline group in the aromatic box were identified by cation-pi contacts with Trp 171, Tyr 210 or Tyr 217 detected as in (51).

### Spectroscopic analysis of MalAzoCh_**β-mer**_

MalAzoCh was synthesized by Enamine Ltd. (Kiev, Ukraine) and compound characterizations (^1^H and ^13^C NMR and HPLC analysis) are described in Figures S1 and S2. MalAzoCh stock solutions were prepared by solubilization of the powder in DMSO at 10mM concentration. They were then divided in aliquots and stocked at -20°C until the day of the experiment.

MalAzoCh solutions were prepared at a concentration of 50 μM in Ringer’s buffer sand irradiated for 15 min at each wavelength with an LED light source (pE-4000, CoolLED). To characterize and quantify the *cis* and *trans* configurations by HPLC–MS, the maleimide group was first reacted with a 10-fold molar excess of β-mercaptoethanol to mimic the conjugation process used for MalAzoCh-C4. β-mercaptoethanol was selected to avoid introducing additional stereogenic center. The thiol-addition was performed at room temperature, and all tubes were protected from ambient light with aluminum foil. All samples were then irradiated at 365 nm, 385 nm or 525 nm for 5 min and subsequently analyzed by HPLC-MS using an Ultimate 3000 RS HPLC coupled with a UV diode array detector and with a Q Exactive mass spectrometer (ThermoFisher Scientific). The MS instrument was equipped with an electrospray ionization source (H-ESI II Probe) and operated in positive mode. Samples were cooled to 4 °C in the autosampler and a 10 µL injection volume was used for each run. Separation was achieved using a Waters ACQUITY UPLC Peptide CSH C18 column (130 Å, 1.7 µm, 50×2.1 mm) at 25°C. Elution was carried out with a mixture of H_2_O (Milli-Q) + 0.1% formic acid (A) and CH_3_CN + 0.1% formic acid (B): 10% B for 0.2 min, 10 to 60% B over 8 min, 60 to 100% B over 0.1 min, and 100% B for 0.6 min (flow rate 0.5 mL/min). Detection was performed at 226 and 361 nm, where the *trans* isomer presents the highest absorbance, and at 316 nm, corresponding to the isosbestic point with the higher sensitivity. Data acquisition and processing were performed using the softwares XCalibur 4.2 and Freestyle 1.7/SP2 (ThermoFisher Scientific). Relative quantification was performed based on the integration of the peak area at 316 nm. MS confirmed the identity of the MalAzoCh_β-mer_ compounds. Representative MS results on sample “Dark” were as follows: C_30_H_42_N_5_O_5_S; m/z 584.2898 (*cis*, Rt 2.86 min) and 584.2896 (*trans*, Rt 3.67 min) ([M]+ calcd 584.2901).

UV-Vis spectra were acquired directly after a 15-min illumination period. For thermal stability assays, the solutions were illuminated and then kept in the dark at room temperature, protected from light with aluminum foil, and absorbance measurements were taken at the time points indicated in the corresponding figures.

### VHH Expression and Purification

The gene encoding the C4 nanobody, containing a C-terminal 6xHis tag and an N-terminal Cys-Ser-Ala (CSA) motif, was cloned into a pFUSE-derived vector (InvivoGen). Expi293F mammalian cells (ThermoFisher) were transfected with the plasmid, and protein expression was performed following the manufacturer’s instructions. The 200 ml cell medium was then loaded on a 5 ml HiTrap TALON crude column (Cytiva) and was purified by affinity chromatography. The column was washed with 5 column volumes of PBS containing 150 mM NaCl, and bound protein was eluted with PBS supplemented with 500 mM Imidazole at pH 7.4. Eluted fractions were then concentrated using Amicon Ultra centrifugal filters 3K-CutOff with regenerated cellulose (Sigma) and further purified by size exclusion chromatography on a Superdex 75 Increase 10/300GL pre-packed column (Cytiva) equilibrated in PBS 150 mM NaCl buffer. Protein identity and purity were assessed by SDS-PAGE and mass spectrometry. The yield of the production was > 7 mg per 200 mL of culture.

### MalAzoCh-C4 conjugation

To reduce the mono-cysteinylated C4 formed during expression, purified CSA-C4 nanobody (25-75 µM) was incubated with a 10-fold molar excess of tris(2-carboxyethyl) phosphine (TCEP) (Sigma). Conjugation was then performed by adding a 10-fold molar excess of MalAzoCh (10 mM in DMSO). The reduction and conjugation reactions were carried out for at least 3 h each, at 4°C under gentle agitation in a ThermoMixerC (Eppendorf). Reaction progress was monitored by HPLC-MS at each step to confirm complete reduction of CSA-C4 and full labeling with MalAzoCh. The excess TCEP and MalAzoCh were removed using a PD MidiTrap desalting column eluted with Ringer’s buffer (Cytiva). MalAzoCh-C4 was used without further concentration. For storage, it was frozen at -20°C directly in the buffer and present no noticeable degradation up to one year.

Analyses were performed using the same equipment described above, with a Waters XBridge BEH C18 column (300 Å, 3.5µm, 100 × 2.1 mm) at 50°C. Samples were cooled to 4 °C in the autosampler and a 2 µL injection volume was used for each run. Elution was conducted with a mixture of H_2_O (Milli-Q) + 0.1% formic acid (A) and CH_3_CN + 0.1% formic acid (B): 10% B for 3 min, 10 to 100% B over 10 min, and 100% B for 7 min (flow rate 0.35 mL/min). Detection was performed at 215 nm, 230 nm and 280 nm. Instrument control and data acquisition were carried out using XCalibur 4.2 (ThermoFisher Scientific). Spectra were deconvoluted using BioPharma Finder 3.2 (ThermoFisher Scientific) with the Intact Mass workflow, employing the ReSpect algorithm. The resulting average masses were compared with theoretical values calculated from the molecular formulas. The same protocol was used for the C4-CSA conjugate.

### Oocyte treatment and injection

Defolliculated, Stage VI *Xenopus* laevis oocytes were purchased from EcoCyte Bioscience, Germany and kept in Barth’s solution (87.34 mM NaCl, 0.66 mM NaNO_3_, 0.72 mM CaCl_2_, 0.82 mM MgSO_4_, 2.4 mM NaHCO_3_, 10 mM HEPES, pH 7.6). Fragments of ovary bags were also obtained from TEFOR Paris-Salay (TPS), France, prepared as previously described (52). For expression, 5-8 ng of cDNA encoding either human α7 nAChR or α7 and β2 nAChR subunits (1:1 ratio), cloned into the pRK5 vector, were injected into the oocyte nucleus using a compressed air injection system. Approximately 1 ng cDNA encoding GFP was co-injected as a reporter gene for successful intracellular injection. Injected oocytes were placed in a 96-well microtiter plate and were incubated in Barth’s solutions at 19 °C for 24-72 h. Injection pipettes were prepared using glass capillaries Drummond Scientific (3-00-203-G/X, WPI), using a Narishige TM PC-100 vertical puller, with temperature parameters set to 69.5°C.

### Two-Electrode Voltage Clamp recordings

TEVC recordings were performed on *Xenopus* laevis oocytes expressing α7 or α7β2 nAChRs, 1-3 days after cDNA injection. Ringer’s buffer (100 mM NaCl, 2.5 mM KCl, 10 mM HEPES, 2 mM CaCl_2_, 1 mM MgCl_2_, pH 7.6) was perfused continuously during the recordings. Currents were recorded using a Clamp Amplifier OC-725D (Warner Instruments), digitized with a Digidata 1550 A and acquired with Clampex 10 (Molecular Devices). Currents were recorded at a holding potential of -60 mV, sampled at 500 Hz and filtered at 100 Hz. Recording pipettes were prepared from Borosilicate glass capillaries with Filament (BF-150-110-7.5, WPI) using a NarishigeTM PC-100 vertical puller, with temperature parameters set at 60.5°C and 49.5°C.

For the illumination protocol, MalAzoCh and MalAzoCh-C4 solutions were pre-illuminated for 15 min with 365, 385 or 525 nm light (LED pE-4000 CoolLED, with powers of 200, 300 and 20 mW, respectively) and were then transferred to the perfusion system of the TEVC set-up, protected from light with aluminum foil. Recordings were performed within 20 min after illumination, to minimize thermal relaxation. Before each perfusion, solutions were briefly re-illuminated. During perfusion, oocytes were illuminated with the corresponding LED wavelength. All experiments were performed at room temperature, in the dark and repeated on at least two independent batches of oocytes. MalAzoCh was prepared as a 10 mM stock solution in DMSO, stored in the dark at -80 °C, and diluted into Ringer’s solution to a final DMSO concentration ≤0.2%.

To calculate the %Activation induced by the free MalAzoCh, the following formula was used:

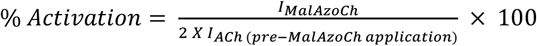

A factor of 2 is applied because the ACh-evoked current measured prior to MalAzoCh application corresponded approximately to the EC_50_.

For MalAzoCh-C4 dose-response experiments, each concentration was tested on a separate oocyte, as full recovery of the current after washout was not always consistently achieved, particularly at higher concentrations Peak current amplitudes were measured for analysis. Data were analyzed with ClampFit (Molecular Devices) and statistical analyses were performed using a Student’s t-test in GraphPad prism. Dose-response curves were fitted to the Hill equation using GraphPad Prism, and data are presented as mean ± standard deviation (SD). Rise and decay time constants were determined in Clampfit by monoexponential fitting of current onset and offset phases.

### Animals

Wild-type C57BL/6J mice (postnatal day 17–27) were obtained from Janvier Laboratories (France). All procedures were carried out in compliance with the European Commission directives 2010/63 on the protection of animals used for scientific purposes and approved by PSL.

### Hippocampal slices preparation

Mice were anesthetized with isoflurane and euthanized by decapitation. Brains were immediately removed and immersed in ice-cold, oxygenated (95% O_2_ / 5% CO_2_) cutting solution containing (in mM): 125 NaCl, 2.5 KCl, 1.25 NaH_2_PO_4_, 26 NaHCO_3_, 10 D-glucose, 15 sucrose, 2 CaCl_2_, 1 MgCl_2_, and 1.06 kynurenic acid.

Coronal slices (250 μm) containing the hippocampus were prepared using a Compresstome (VF-200, Precisionary Instruments, USA), and incubated in the same oxygenated cutting solution at 35 °C for 8 minutes. Following incubation, slices were transferred to a holding chamber containing room temperature, oxygenated recording aCSF composed of (in mM): 125 NaCl, 2.5 KCl, 1.25 NaH_2_PO_4_, 26 NaHCO_3_, 10 D-glucose, 15 sucrose, 2 CaCl_2_, and 1 MgCl_2_. Slices were stored in this solution for the duration of the experimental day.

### Whole-cell patch-clamp recordings in CA1 interneurons

Individual slices were transferred to a recording chamber and continuously perfused with oxygenated aCSF at room temperature and a rate of 2 mL/min. Whole-cell patch-clamp recordings were performed using an Axopatch 200B amplifier connected to a Digidata 1550 Low-Noise acquisition system (Molecular Devices, USA) and controlled via Clampex (pClamp 10.5, Molecular Devices).

Signals were low-pass filtered at 2 kHz (Bessel filter) and digitized at 10 kHz. Recordings were extracted using Clampfit (Molecular Devices) and analyzed in R. Patch pipettes were pulled from G150TF-3 borosilicate glass capillaries (Warner Instruments, USA) using a horizontal puller (P-87, Sutter Instruments, USA), with a final resistance of 5–8 MΩ. Cells were visualized using an upright microscope equipped with Dodt contrast optics and illuminated with a white light source (Scientifica, UK). The internal pipette solution contained (in mM): 135 K-gluconate, 10 HEPES, 5 KCl, 2 MgCl_2_, 0.1 EGTA, 2 Mg-ATP, and 0.2 Na-GTP. The pH was adjusted to 7.35 with 8 N KOH, and the osmolarity was set to 290–300 mOsm. Voltage- and current-clamp recordings were performed on hippocampal interneurons located in the stratum radiatum and stratum lacunosum of the CA1 region. Once a cell was patched and held at ™60 mV, baseline noise was recorded prior to compound application. A cell was classified as responding if the evoked current amplitude exceeded three times the standard deviation of the baseline noise. A puff pipette containing 3 µM MalAzoCh-C4 diluted in aCSF was positioned in the same focal plane, approximately 20 µm from the recorded cell, using micromanipulators (Scientifica, USA). Compound application was achieved using a pneumatic picopump (PV-800, World Precision Instruments, USA) connected to a pressurized air system, delivering MalAzoCh-C4 (3 µM) at 2-6 psi. Optical stimulation was delivered via the microscope using two LED light sources (pE-2, CoolLED) at 380 nm and 525 nm wavelengths. The measured light power at the objective output was 6.5 and 15 mW, corresponding to an intensity of approximately 5 and 12 mW/mm^2^ at the focal plane, respectively.

### Statistical analysis for the patch-clamp recordings

Sample sizes were not determined using statistical power calculations. Data are presented as mean ± standard error of the mean (SEM). The number of observations(n) for each condition, along with the statistical tests used, are specified in the corresponding figures and/or figure legends. Unless otherwise noted, group comparisons were made using non-parametric tests (Wilcoxon or Friedman), as data did not meet the assumptions of normality based on the Shapiro–Wilk test (p > 0.05). A p-value greater than 0.05 was considered not statistically significant.

## ASSOCIATED CONTENT

### Supporting Information

The following file is available free of charge. Supplemental figures

Details of chemical synthesis, biochemistry, mass characterization, additional electrophysiological data from oocytes and brain tissue complementing the main text. Tables with electrophysiological values, kinetic values and mass spectrometry values. (PDF)

## AUTHOR INFORMATION

### Author Contributions

Conceptualization: AM, PJC

Methodology: MV, KS, SB, GA, PF, LP, AB, AM, PJC

Investigation: MV, SB, KM, GA (nanobody production and conjugation), MV (TEVC), KS (patch-clamp electrophysiology), AB (in silico docking)

Supervision: AM, PJC

Writing: MV, SB, KS, AB, AM, PJC

## ACKNOWLEDGMENT

The authors would like to thank the animal facility at ESPCI, Paris, and the DIM C-BRAINS (Région Ile-de-France) and NeuroPSL (PSL University) for their support.

## ABBREVIATIONS

BBB: blood-brain barrier
ECD: extracellular domain
MLA: methyllycaconitine
MS: mass spectrometry
nAChR: nicotinic acetylcholine receptor
NAM: negative allosteric modulator
PAM: positive allosteric modulator
PLC: photochromic ligand
PORTL: photochromic orthogonal remotely tethered ligand
PTL: photochromic tethered ligand
SAL: silent allosteric ligand
TCEP: tris(2-carboxyethyl) phosphine
TEVC: two-electrode voltage clamp
TMD: transmembrane domain.

## Insert Table of Contents artwork here

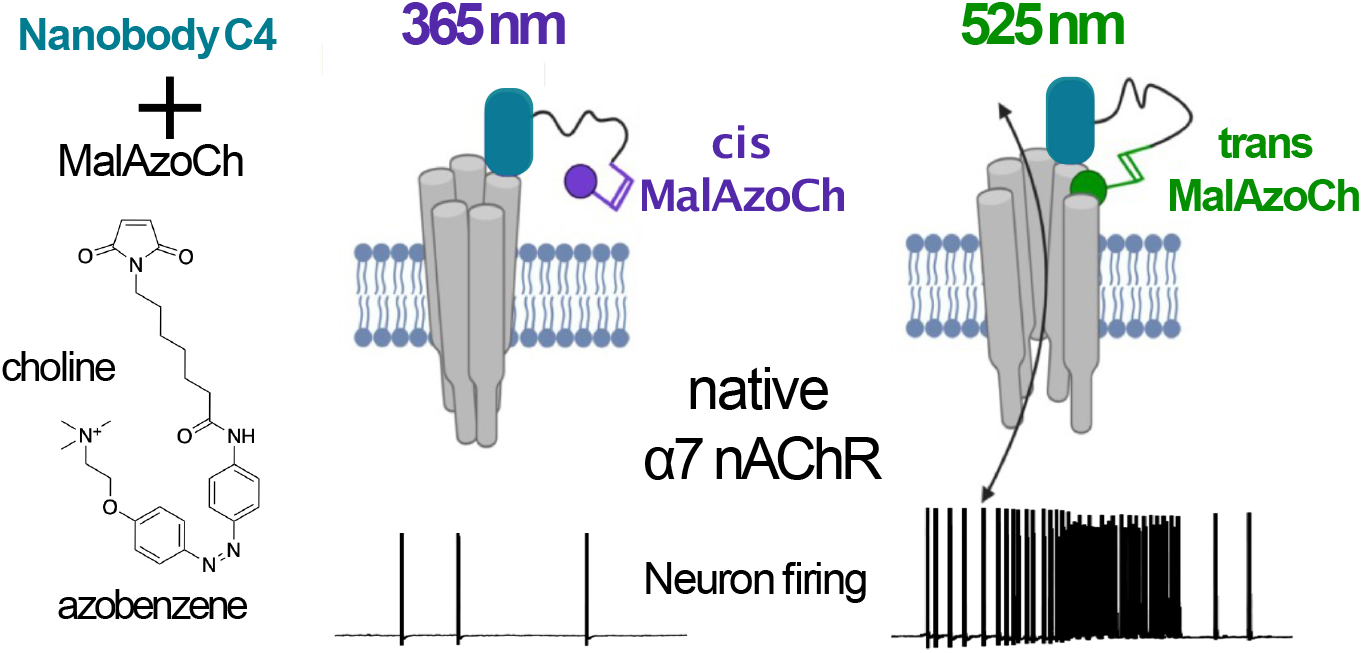

