## supplementary file for "AN OPTONANOBODY FOR REVERSIBLE PHOTOACTIVATION OF RECOMBINANT AND NATIVE α7 NICOTINIC RECEPTORS"

**TABLE OF CONTENTS**


| | |
|---|---|
| Figure S1 | 3 |
| Figure S2 | 4 |
| Figure S3 | 5 |
| Figure S4 | 6 |
| Figure S5 | 7 |
| Figure S6 | 8 |
| Figure S7 | 9 |
| Figure S8 | 10 |
| Figure S9 | 11 |
| Figure S10 | 12 |
| Figure S11 | 13 |
| Figure S12 | 14 |
| Table S1 | 15 |
| Table S2 | 16 |
| Table S3 | 17 |

**A**

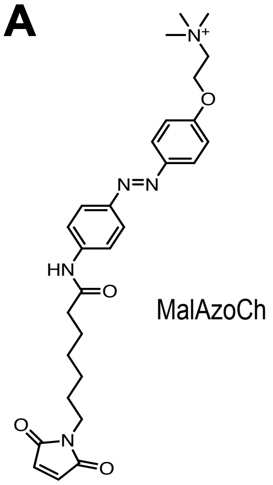


**B**

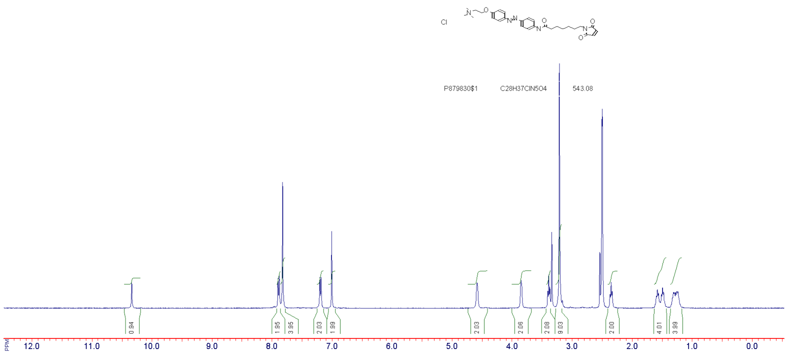


**C**

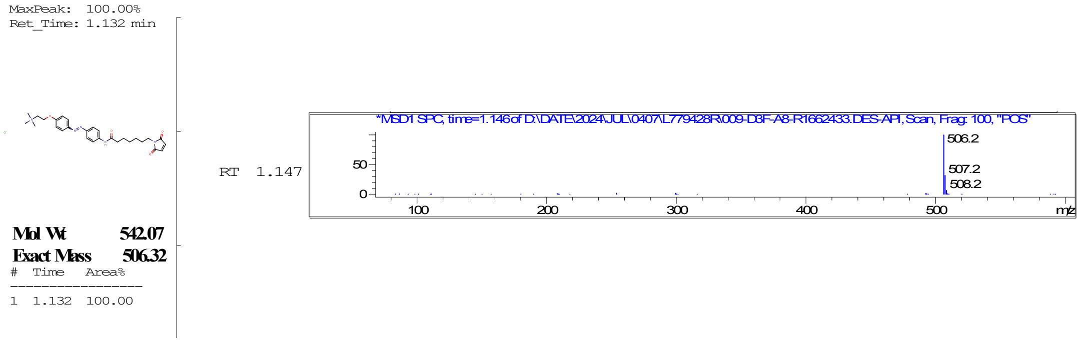


**Fig. S1. Chemical characterization of MalAzoCh. (A)** Structural formula of MalAzoCh. (**B**) $^{1}$H NMR spectrum of the *trans* MalAzoCh in the dark solubilized in DMSO and (**C**) HPLC-MS analysis of MalAzoCh. The peak observed at retention time 1.147 shows the expected mass profile of MalAzoCh.

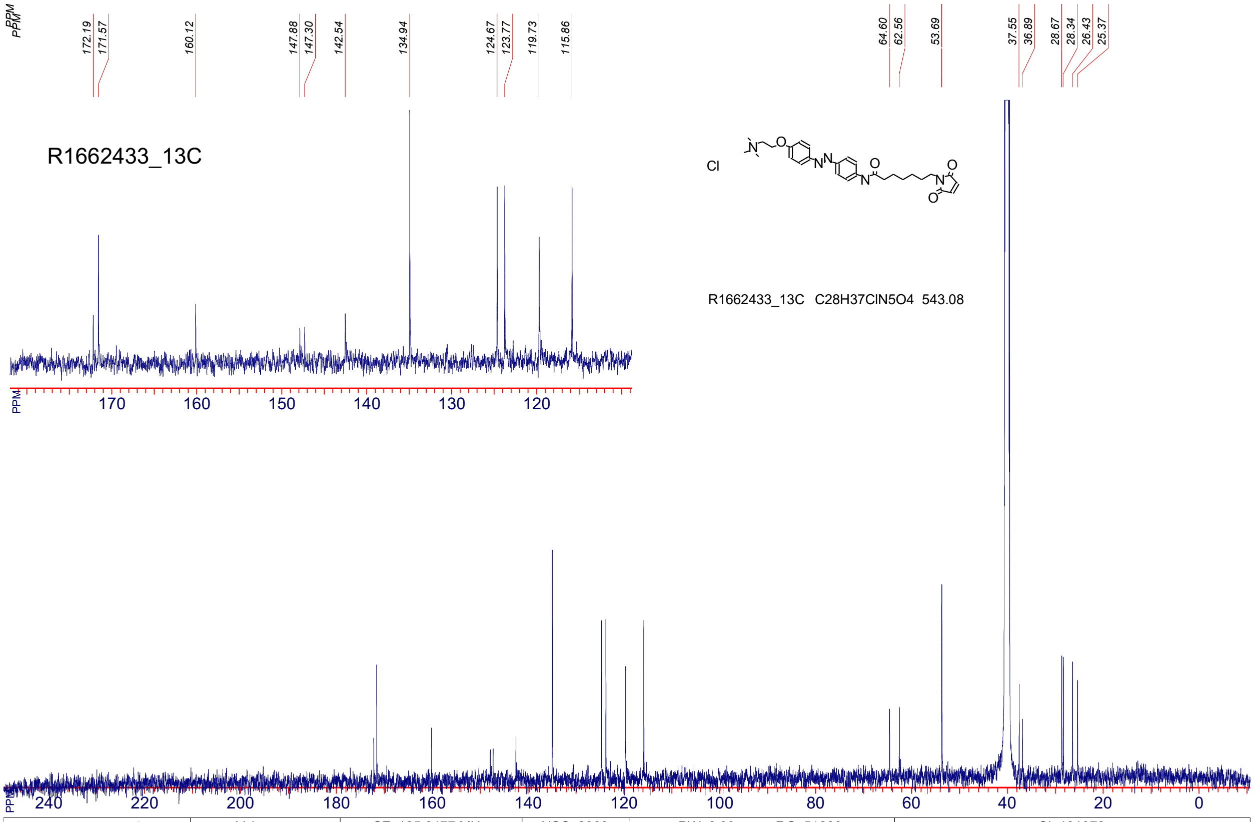


| R1662433_13C | Vybornov | SF: 125.6177 MHz | NSC: 2389 | PW: 0.00 usec, RG: 51200 | SI: 131072 |
|---|---|---|---|---|---|
| Date: 24-May-2026 | Solvent: DMSO | SW: 32680 Hz | TE: 683 K | AQ: 1.57 sec, RD: 0.00 sec | |

**Fig. S2. $^{13}$C NMR spectrum of MalAzoCh solubilized in DMSO.**

**A**

**C4CSA** MGSSHHHHHHAAAAQVQLVESGGGLVQAGGSLKLSCAASGFTFAHYAMVWFRQAPGKERE
FVAGISWSGASTYYASSVKGRFTISRDNAKNTVYLQMNSLKPEDTAVYYVAAARFGVGVD
DDYSYWGQGTQVTVSSGGGGSGGGGSGGGGS**CSA**

**CSAC4** MG**CSA**GGGGSGGGGSQVQLVESGGGLVQAGGSLKLSCAASGFTFAHYAMVWFRQAPGKER
EFVAGISWSGASTYYASSVKGRFTISRDNAKNTVYLQMNSLKPEDTAVYYVAAARFGVGV
DDDYSYWGQGTQVTVSSAAAHHHHHHGS*

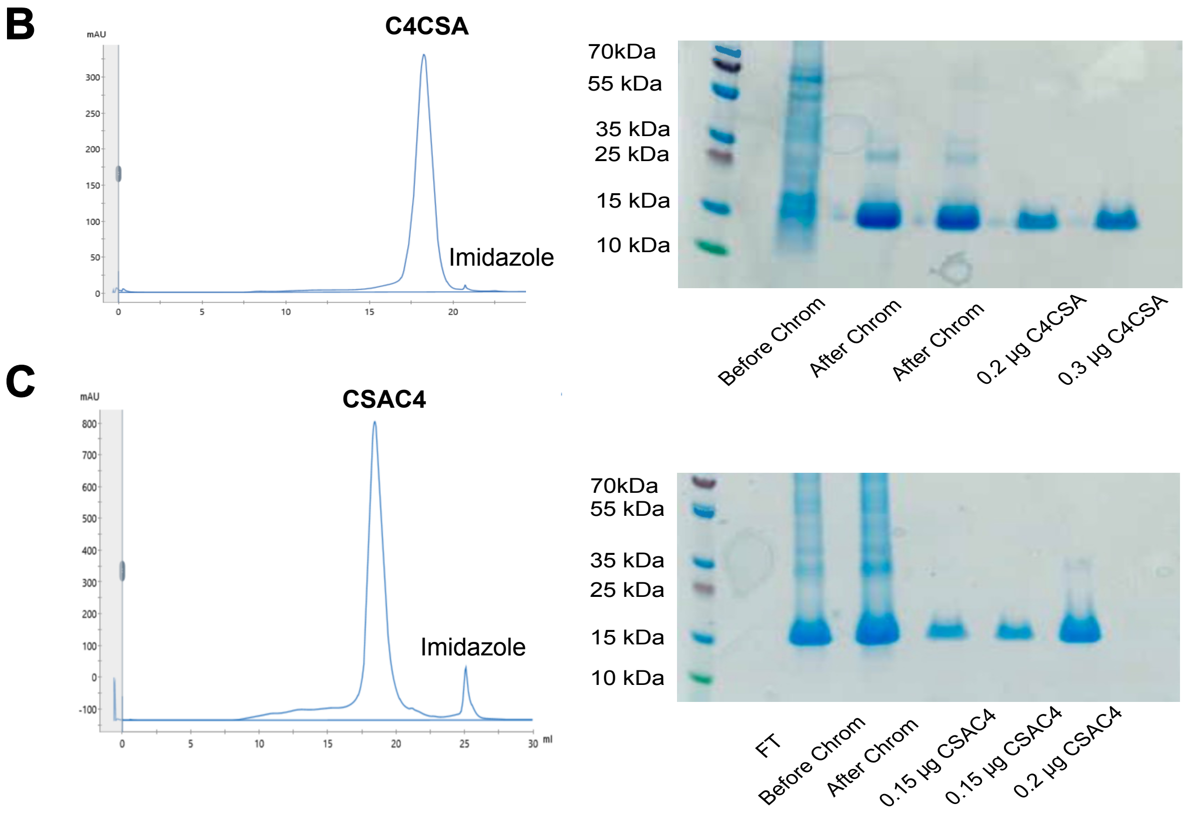


**Fig. S3. Production and purification of the C4 constructs. (A)** Sequence of the two C4 constructs, C4CSA and CSAC4 used in this study showing the His-Tag in red and the CSA moiety for conjugation in bold. **(B)** and **(C)** Left panel: Size-exclusion chromatography profiles of the C4 constructs and right panel: SDS PAGE analysis of the C4 constructs. The SDS-PAGE gel displays the flowthrough (FT) following protein concentration, as well as the protein profiles before and after affinity chromatography, in addition to the different fractions obtained from size-exclusion chromatography (SEC) of the C4 constructs.

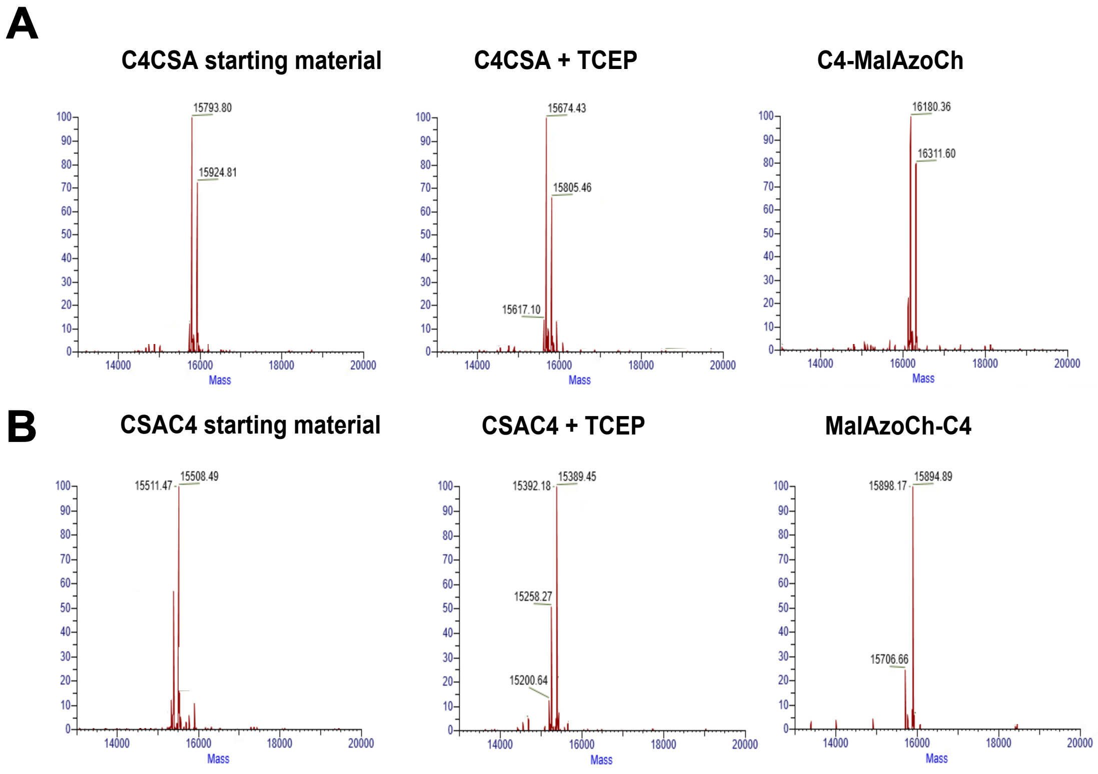


**Fig. S4. MS analyses of compounds involved in MalAzoCh conjugation to C4.** MS deconvoluted spectra of the two C4 constructs, showing, from left to right: the C4 cysteinylated starting material which may result from either a post-translational modification of the protein or a cysteinylation in the expression medium, the reduced C4 following reaction with TCEP and the final conjugate C4 with AzoCh tethered at the C-terminal (A) and at the N-terminal (B) region. The deconvoluted spectra demonstrate that the conjugation reactions are complete. Expected Mr are indicated in Table S1 with the corresponding assignments of the observed compounds.

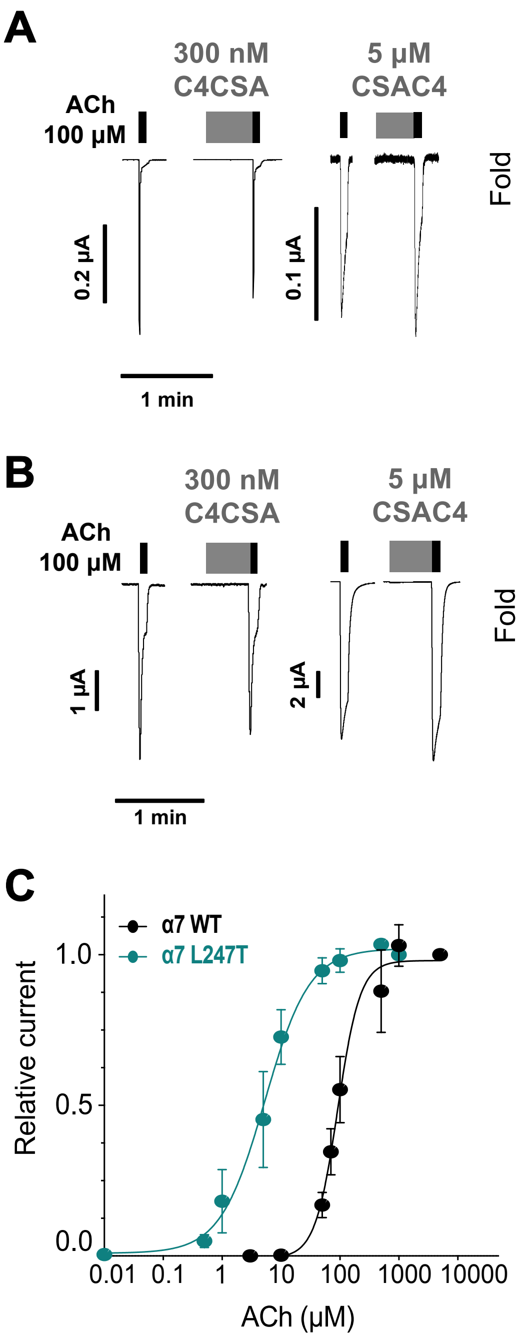


**Fig. S5. Electrophysiological analysis of the C4 constructs on α7 WT (A) and L247T (B) nAChR.** The C4 constructs were applied for a 30-second duration, followed by an immediate ACh-application to assess any potential modulatory effects. The right panel shows the fold ACh-currents recorded after the C4 application. The results demonstrate that the C4 constructs are silent allosteric ligand. Values were submitted to an unpaired *t*-test with *ns*, $p > 0.05$ and **C.** ACh dose-response curves of α7 WT and L247T with mean ±SD ($n$ = 4-6).

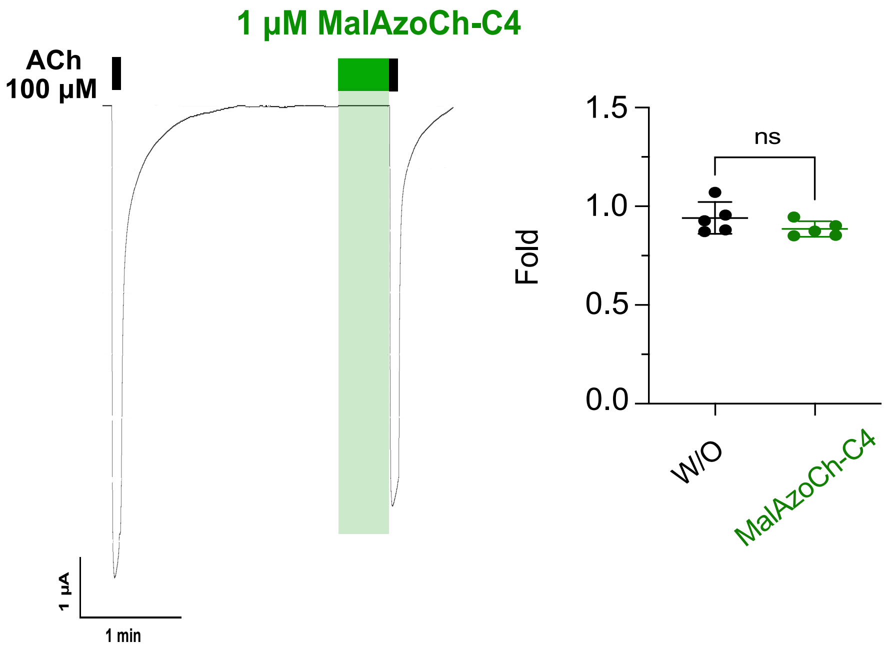


**Fig. S6. Electrophysiological analysis of 1 µM MalAzoCh-C4 on α4β2 nAChR.** MalAzoCh-C4 was applied for a 30-second duration, followed by an immediate ACh-application to assess any potential modulatory effects. The right panel shows the fold ACh-current remaining without (W/O) and after MalAzoCh-C4 application. The results demonstrate that MalAzoCh-C4 does not induce any modulatory effects on α4β2 nAChR. Values were submitted to an unpaired *t*-test with *ns*, $p > 0.05$.

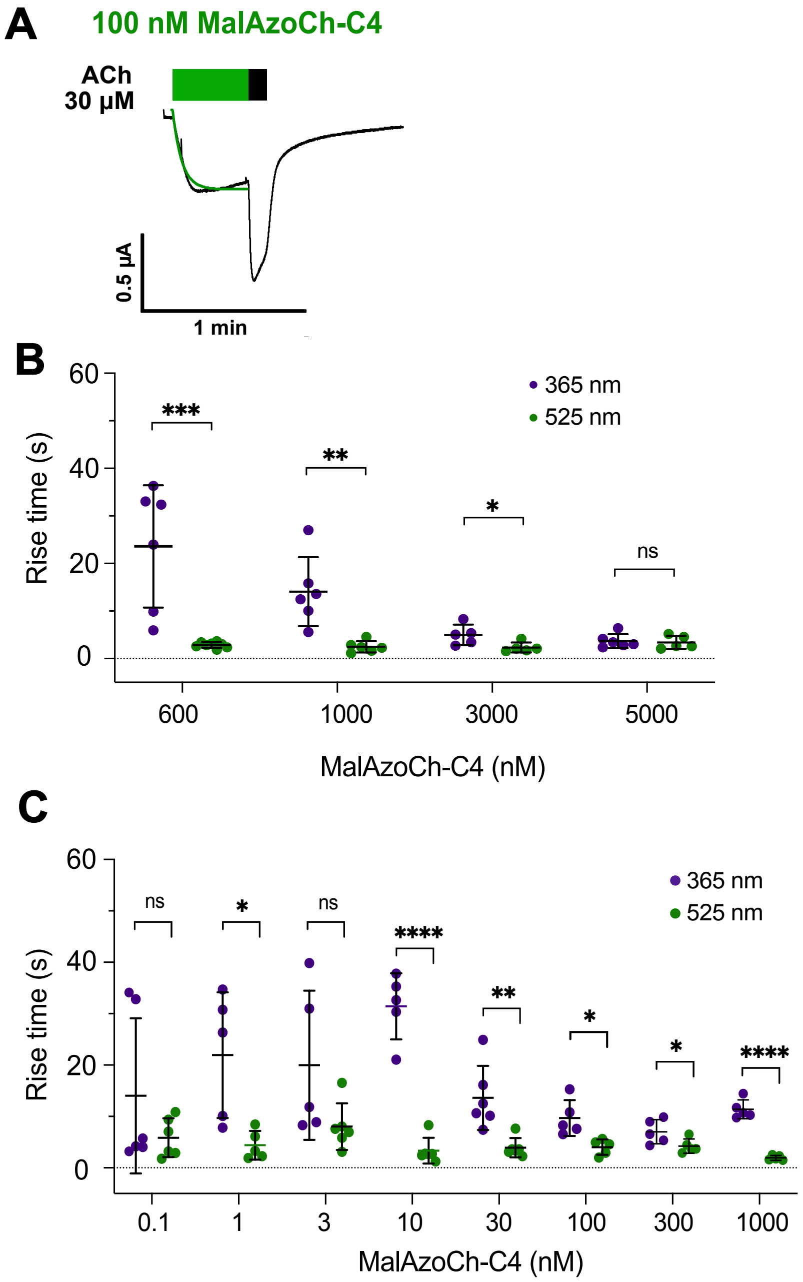


**Fig. S7. Activation kinetics of MalAzoCh-C4 at 365 nm and 525 nm illumination. (A)** Example of a single exponential fitting of current trace rise time and corresponding values on **(B)** α7 WT and **(C)** α7 L247T. Values were submitted to an unpaired *t*-test with *ns*, $p > 0.05$, *, $p < 0.05$, **, $p < 0.001$, ****, $p < 0.0001$. Bars are mean ± SD.

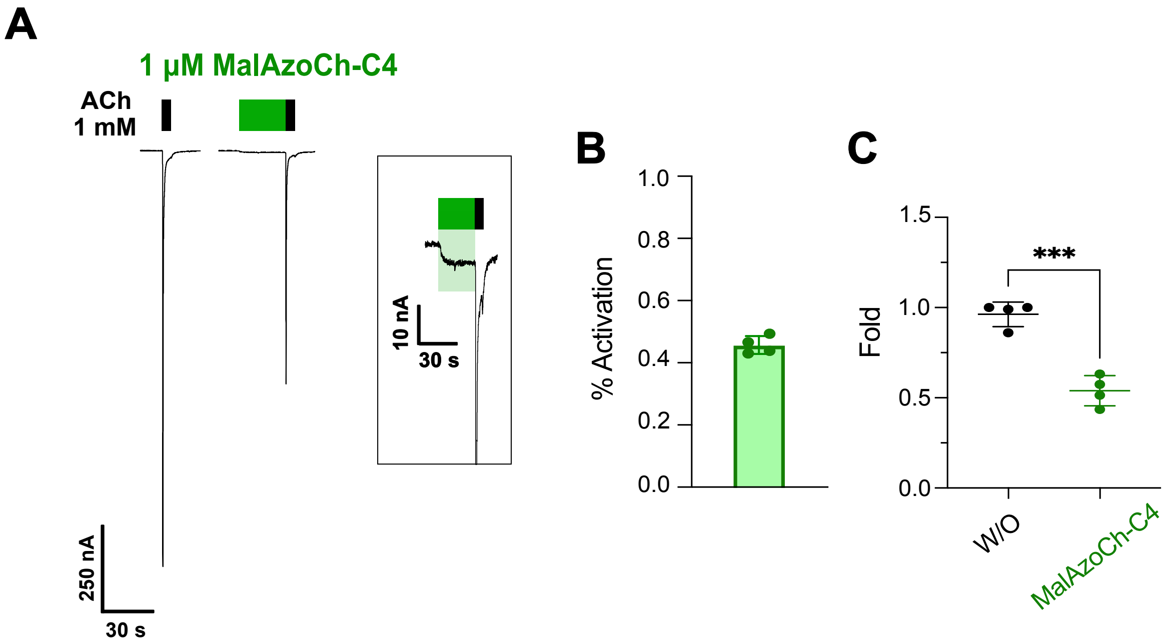


**Fig. S8. MalAzoCh-C4 is an agonist of α7β2 nAChR. (A)** Current traces of oocytes expressing α7β2 nAChR ($n$ = 4). 1 µM of MalAzoCh-C4 was applied for 30 seconds under green light and was shown to activate and inhibit significantly post-perfusion ACh-induced current. **(B)** Bar graph showing the relative activation of 1 µM of MalAzoCh-C4 normalized to the ACh-current. **(C)** Fold current remaining after MalAzoCh-C4 application. Values were submitted to an unpaired $t$-test with ***, $p < 0.001$. Bars are mean ± SD.

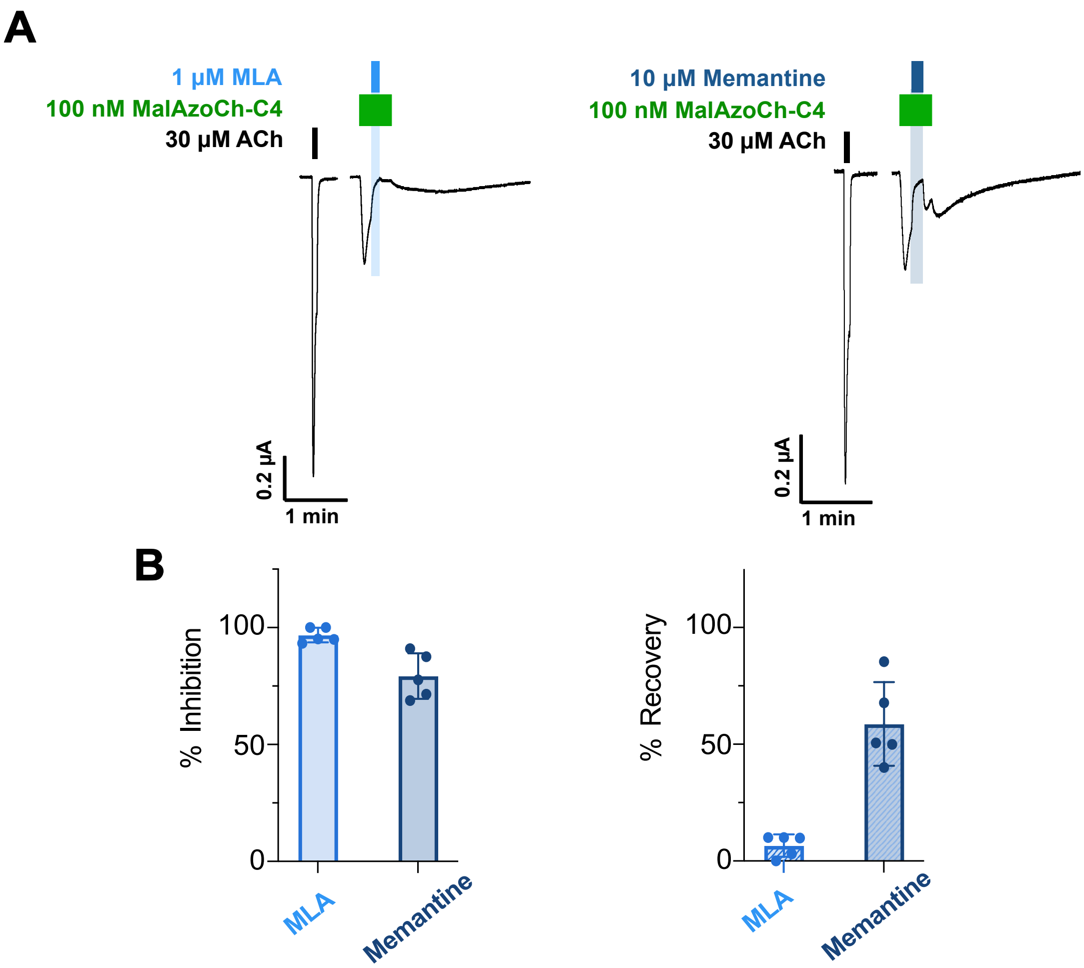


**Fig. S9. Inhibition of MalAzoCh-C4. (A)** Current traces showing successive application of 100 nM MalAzoCh-C4 alone, 100 nM MalAzoCh-C4 co-applied with 1 µM MLA (left panel) or 10 µM memantine (right panel), and then 100 nM MalAzoCh-C4 alone on α7 L247T. **(B)** Bar graph summarizing the percentage of inhibition and recovery following the application and subsequent removal of the inhibitor molecules. Bars are mean ± SD.

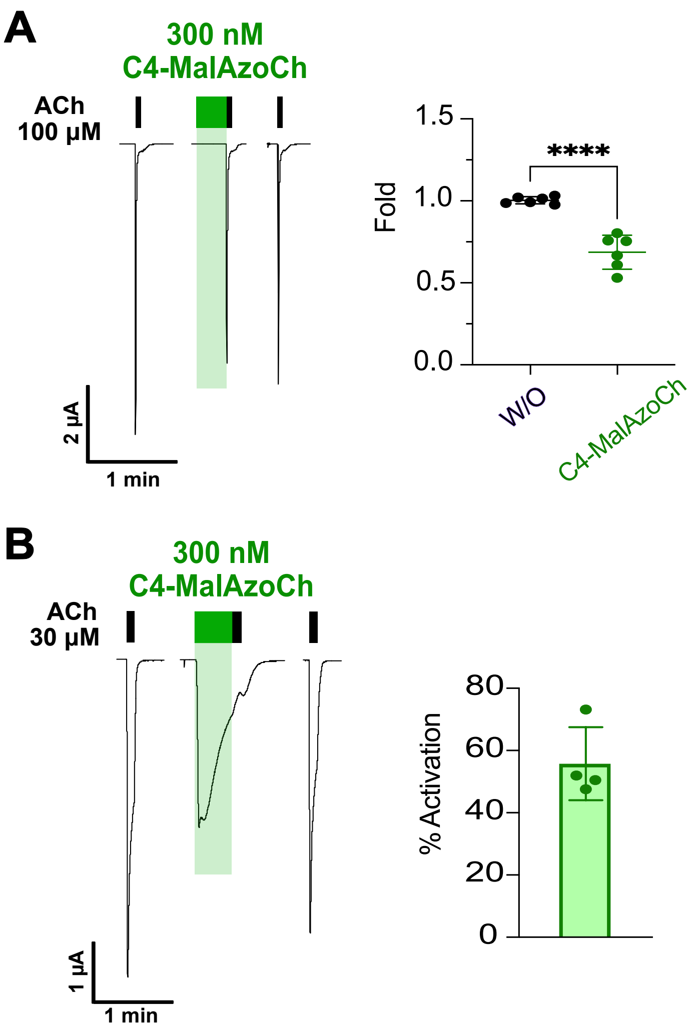


**Fig. S10. C4-MalAzoCh acts as an agonist of α7 nAChR. (A)** Current traces of oocytes expressing α7 nAChR (*n* = 6). 300 nM of C4-MalAzoCh were applied for 30 seconds under green light and elicit no significant current but decrease the amplitude of the ACh post-perfusion application. The fold current remaining is shown on the right panel. Values were submitted to an unpaired *t*-test with ****, $p < 0.0001$ and **(B)** Current traces of oocytes expressing α7 L247T (*n* = 4) showing robust activation by 300 nM C4-MalAzoCh. Bar graph shows corresponding activation as compared to ACh maximal currents. Bars are mean ± SD.

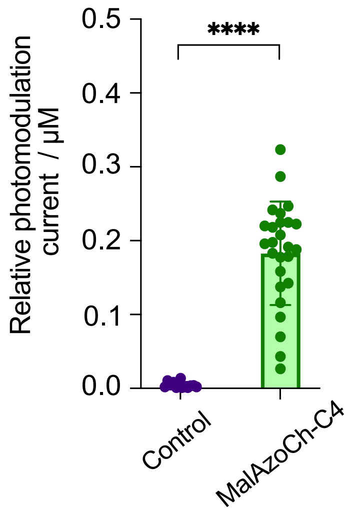


**Fig. S11. Photomodulation of α7 L247T with 30 µM ACh 10 nM MalAzoCh-C4.** Graph illustrating the normalized photomodulation ratio $I_{525}/I_{365}$ of current, normalized to the 30 µM ACh response for each oocyte expressing α7 L247T ($n$ = 6). The green bar represents the photomodulatio ratio $I_{525}/I_{365}$ for 10 nM MalAzoCh-C4 and the black corresponds to the control experiment, where oocytes expressing α7 L247T were subjected to a prolonged application of 30 µM ACh with continuous light photoswitching (365 nm and 525 nm). This corresponds to the negative control. ****, $p < 0.0001$, Mann-Whitney test.

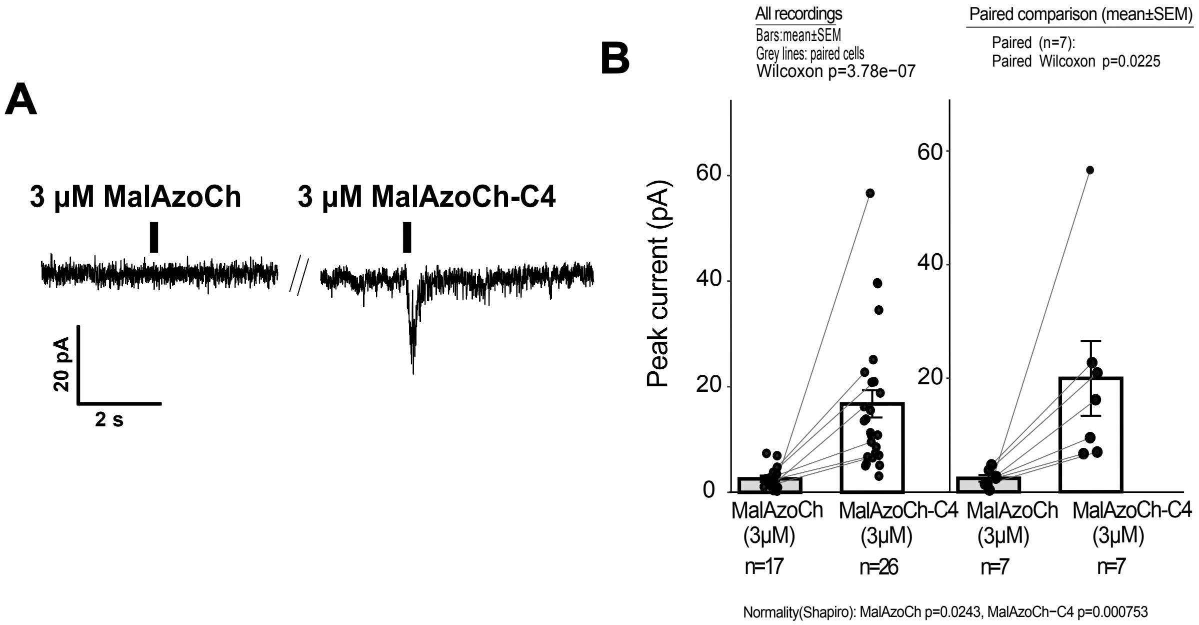


**Fig. S12. Conjugation to C4 is required for MalAzoCh (3 µM) to activate native α7 nAChRs. (A)** Representative whole-cell voltage-clamp recordings (−60 mV) from hippocampal interneurons showing inward currents evoked by a 200 ms puff application of MalAzoCh (3 µM) or MalAzoCh-C4 (3 µM). **(B)** Summary of peak current amplitudes. A cell was classified as responding when the evoked current amplitude exceeded three times the standard deviation of the baseline noise. MalAzoCh-C4 elicited currents in 100% of the recorded cells (n = 26 cells from 12 mice) whereas MalAzoCh never elicited current (n = 17 cells from 3 mice; unpaired Wilcoxon test, $p = 3.78 \times 10^{-7}$). Cells that received both compounds are connected by lines (n = 7 cells from 2 mice; paired Wilcoxon test, p = 0.022). Data are shown as mean ± SEM.

| | C4CSA | | | CSAC4 | | |
|---|---|---|---|---|---|---|
| | Cysteinylated starting material | Reduced (+TCEP) | C4-MalAzoCh | Cysteinylated starting material | Reduced (+TCEP) | MalAzoCh-C4 |
| **Full sequence with Met** | 15925.25 | 15806.11 | 16312.73 | 15508.87 | 15389.73 | 15896.34 |
| **-Met** | 15794.06 | 15674.92 | 16181.53 | 15377.67 | 15258.53 | 15765.14 |
| **-Met-Gly** | | | | 15320.62 | 15201.48 | 15708.09 |

**Table S1. Expected $M_r$ are of compounds involved in MalAzoCh conjugation to C4.** The corresponding mass spectra are shown in Fig S3.

| | $EC_{50}$ (µM) | $IC_{50}$ (nM) | nHill |
|---|---|---|---|
| α7 WT / ACh | 108.5 ± 30.12 | | 1.94 ± 0.74 |
| α7 WT / MalAzoCh-C4 (365 nm) | | 1197 ± 222.7 | 11.47 ± 10.12 |
| **α7 WT / MalAzoCh-C4 (*525 nm*)** | | 574 ± 92.2 | 2.13 ± 0.52 |
| α7 L247T / ACh | 5.4 ± 2.5 | | 1.43 ± 0.52 |
| α7 L247T / MalAzoCh-C4 (*365 nm*) | | 606.5 ± 320.7 | 5.45 ± 6.01 |
| α7 L247T / MalAzoCh-C4 (*525 nm*) | | 26.1 ± 21.1 | 0.83 ± 0.36 |

**Table S2. $EC_{50}$ and $IC_{50}$ values for current responses to ACh and MalAzoCh-C4.**

| α7 WT | | | | |
|---|---|---|---|---|
| | 365 nm | | 525 nm | |
| MalAzoCh-C4 (nM) | Rise Time (s) | SD | Rise Time (s) | SD |
| 600 | 23,59 | 12,88 | 2,86 | 0,59 |
| 1000 | 14,07 | 7,23 | 2,48 | 1,19 |
| 3000 | 4,97 | 2,17 | 2,32 | 1,05 |
| 5000 | 3,69 | 1,45 | 3,40 | 1,37 |
| α7 L147T | | | | |
| MalAzoCh-C4 (nM) | Rise Time (s) | SD | Rise Time (s) | SD |
| 0,1 | 14,02 | 15,09 | 5,87 | 3,78 |
| 1 | 21,95 | 12,24 | 4,41 | 2,80 |
| 3 | 19,97 | 14,50 | 8,02 | 4,52 |
| 10 | 31,42 | 6,44 | 3,36 | 2,50 |
| 30 | 13,63 | 6,24 | 3,94 | 1,88 |
| 100 | 9,71 | 3,48 | 4,03 | 1,50 |
| 300 | 7,02 | 2,35 | 4,24 | 1,39 |
| 1000 | 11,39 | 1,83 | 1,94 | 0,46 |
| α7 WT | Rise Time (ms) | SD | | |
| ACh (100 µM) | 782 | 462 | | |

**Table S3. Rise time values upon MalAzoCh-C4 application on α7 WT and L247T.**
